# Single-molecule chromatin configurations link transcription factor binding to expression in human cells

**DOI:** 10.1101/2024.02.02.578660

**Authors:** Benjamin R. Doughty, Michaela M. Hinks, Julia M. Schaepe, Georgi K. Marinov, Abby R. Thurm, Carolina Rios-Martinez, Benjamin E. Parks, Yingxuan Tan, Emil Marklund, Danilo Dubocanin, Lacramioara Bintu, William J. Greenleaf

## Abstract

The binding of multiple transcription factors (TFs) to genomic enhancers activates gene expression in mammalian cells. However, the molecular details that link enhancer sequence to TF binding, promoter state, and gene expression levels remain opaque. We applied single-molecule footprinting (SMF) to measure the simultaneous occupancy of TFs, nucleosomes, and components of the transcription machinery on engineered enhancer/promoter constructs with variable numbers of TF binding sites for both a synthetic and an endogenous TF. We find that activation domains enhance a TF’s capacity to compete with nucleosomes for binding to DNA in a BAF-dependent manner, TF binding on nucleosome-free DNA is consistent with independent binding between TFs, and average TF occupancy linearly contributes to promoter activation rates. We also decompose TF strength into separable binding and activation terms, which can be tuned and perturbed independently. Finally, we develop thermodynamic and kinetic models that quantitatively predict both the binding microstates observed at the enhancer and subsequent time-dependent gene expression. This work provides a template for quantitative dissection of distinct contributors to gene activation, including the activity of chromatin remodelers, TF activation domains, chromatin acetylation, TF concentration, TF binding affinity, and TF binding site configuration.

## Introduction

Genomic regulatory elements, such as enhancers and promoters, coordinate the binding and activity of multiple transcription factors (TFs), DNA-sequence-specific binding proteins, to drive transcription in human cells. On average, endogenous enhancers contain binding sites for 5-6 TFs^1^, and in many tractable experimental systems, increasing the number of sites leads to strongly nonlinear effects on gene expression^2,3^. Many hypotheses have been put forward to explain this observed synergy between TF binding site number and gene expression^4^. Synergy might arise at the level of TF binding, either through direct physical interactions between TFs^5^, from a thermodynamic competition between TFs and nucleosomes for DNA occupancy^6^, or through the effects of cofactors (such as chromatin remodelers or acetyltransferases) which aid in nucleosome eviction^7,8^. Alternatively, synergy could be the result of TFs acting on different steps of the transcription cycle and cooperating to drive super-additive expression once bound^9,10^.

Arbitrating between these distinct molecular mechanisms would be immensely enabled by a capacity to directly measure molecular occupancy configurations at the resolution of the number and position of individual proteins bound to specific regulatory elements. Current methods, such as ChIP-seq or ATAC-seq, provide aggregate and often non-quantitative information on these DNA occupancy states and cannot detail the correlated binding of multiple proteins across a single chromatin fiber. However, single-molecule footprinting methods that use methyltransferases to mark accessible sites^11–15^ can provide quantitative measurements of these binding states across individual molecules. Therefore, these tools have the potential to enable investigation and first-principles modeling of the full “chain of causality” from DNA sequence to intermediate binding configurations at the level of chromatin to resulting gene expression.

To this end, we have developed a platform for investigating the relationship between molecular configurations and resulting gene expression at a synthetic regulatory element to dissect the relationship between TF occupancy and transcriptional output. In two reporter systems engineered into the genome – one with increasing numbers of synthetic TF binding sites and one with increasing numbers of interferon-stimulated response elements – we use single-molecule footprinting methods to measure TF binding events and link them to gene expression both at steady state and over time. We demonstrate that non-linear transcriptional responses to increasing numbers of TF binding sites is a result of the apparent cooperative binding of TFs, which itself is driven by the active eviction of nucleosomes by activation domain-recruited chromatin remodelers. Furthermore, we observe that the capacity of TFs to activate transcription is proportional to the average number of bound TFs, and that different TFs possess distinct capacities (or potencies) once bound. We develop a non-equilibrium steady-state model inspired by partition-function-based statistical mechanics that can predict observed distributions of enhancer binding states. Through drug-based inhibition, we measure the contributions of the chromatin remodeler BAF and acetyltransferase p300 to 1) enabling TF binding at the enhancer and 2) driving states permissive for expression at the promoter, observing that BAF influences both while p300 influences only promoter activity. Finally, we develop a model capable of explaining chromatin state and gene expression over time in both the synthetic and endogenous reporter systems, revealing distinct mechanisms and kinetics of activation in each.

### SMF reveals molecular binding states at promoters and promoter-proximal enhancers

To systematically measure TF binding and assess the effects of this binding on transcription, we created a library of synthetic reporter constructs consisting of promoter-proximal enhancers containing variable numbers of Tet operator (TetO) sequences spaced 21 bp apart in a random sequence context starting 170 bp upstream of a minimal promoter (minCMV) driving reporter gene expression (Citrine). We then integrated these constructs at the AAVS1 safe-harbor locus in K562 cells constitutively expressing the synthetic TF rTetR-VP48, which binds to TetO upon the addition of doxycycline^16^ (dox) (**Fig. 1A**). This synthetic, inducible, perturbable enhancer-promoter system allows us to vary the composition of TF binding sites (e.g. by engineering a variable number of TetO sites into the genome), the activation domains associated with the TF (e.g. by engineering different rTetR fusion proteins, with or without activation domains), and the effective concentration of TF available for DNA binding (e.g. by varying dox concentration) to explore how these variables affect TF occupancy and gene expression.

**Figure 1:**
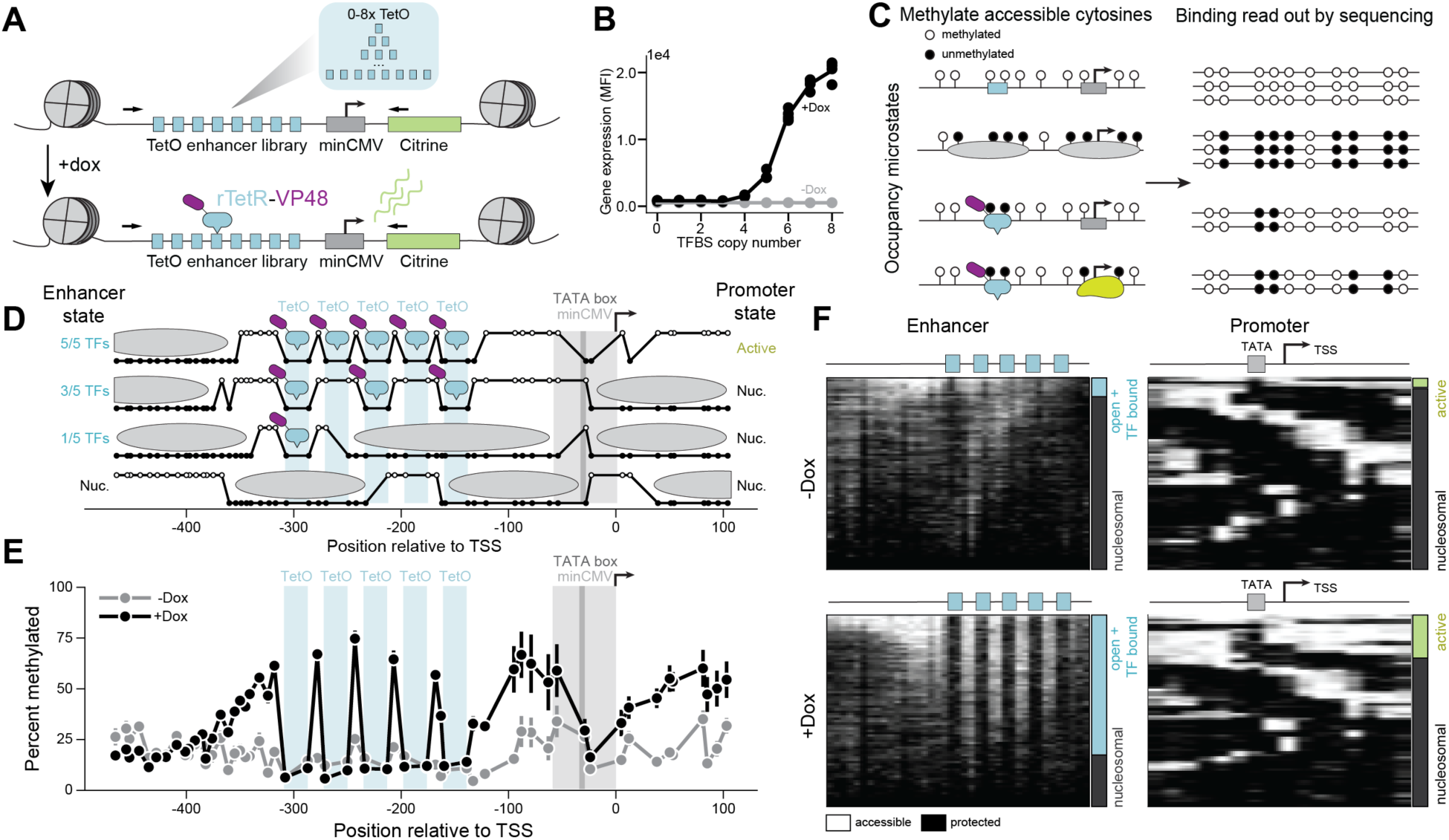
Single-molecule footprinting reveals TF occupancy and promoter state at an engineered, genome-integrated expression reporter system. A) Variable numbers (0-8) of TetO binding sites (blue) upstream of a minCMV (gray) promoter and Citrine (green) gene were engineered into the AAVS1 locus in K562 cells expressing rTetR-VP48. Binding of rTetR-VP48 is inducible upon addition of doxycycline (dox) to cell media. Black arrows are the primers used to sequence the reporter. B) Mean fluorescent intensity (MFI) of the Citrine reporter measured using flow cytometry after 24 hours of 1,000 ng/ml doxycycline exposure (black) as a function of the number of transcription factor TetO binding sites (TFBS). C) Schematic describing the single-molecule footprinting (SMF) assay for determining the molecular state of regulatory elements or promoters. Accessible GpCs are methylated (white circles) while inaccessible GpCs remain unmethylated (black circles). D) Example SMF molecules with 5 TetO sites, along with the molecular configuration interpretations for these molecules. The high state represents a methylated, accessible GpC, while a low state represents an unmethylated, inaccessible GpC. E) Aggregated data obtained for the construct with 5 TetO sites present, with (black) and without (gray) dox present. Error bars represent standard error of the mean from 4 biological replicates. F) Summary plots of single molecules (individual rows) observed by SMF for enhancers (left) and promoters (right) of constructs with 5 TetO sites without (top) and with (bottom) dox induction. Each GpC spans an equal width in this representation; methylated (accessible) Cs in white and unmethylated (protected) Cs in black. The bars at the right of each summary plot show the fraction of enhancer sites with one or more TF bound (open + TF bound; blue) or fully nucleosomal (gray) and the fraction of promoters that are not bound by a nucleosome (active; green) or bound by a nucleosome (gray).

We observed dox-dependent, non-linear increases in gene expression for increasing numbers of TetOs in the enhancer (**Fig. 1B, Fig. S1A**). Gene expression exhibited minimal changes between 24 and 48 hours of dox treatment, demonstrating that the system is at steady state by 24 hours (**Fig. S1B**). To measure the molecular configurations of TFs, nucleosomes, and transcriptional machinery at our synthetic reporter constructs *in vivo*, we adapted and applied a methyltransferase-based single-molecule footprinting (NoME-seq/SMF) assay^11,12,15^. We methylated accessible cytosines in GpC contexts by treating isolated nuclei with the recombinant methyltransferase M.CviPI and measured which cytosines were methylated by enzymatically converting unmethylated cytosines to uracil with a modified EM-seq protocol^17^, amplifying reporter constructs using primers that bind to converted DNA, then sequencing the resulting amplicons (**Fig. 1C**, Methods). In each sequencing read, the conversion status of each GpC cytosine reports the accessibility state at that nucleotide on a single reporter construct molecule in a cell. We flanked each TetO with a pair of GpCs and inserted GpCs every 8 bp on average across the reporter construct to enable high resolution accessibility measurements of each sequence. We confirmed that methylation and enzymatic conversion were efficient (**Fig. S1C-E**), and we optimized lysis conditions, cell number, and methylation time to increase assay sensitivity (**Fig. S1F-G**). We then performed SMF, in tandem with cellular perturbations (described below), and called binding and nucleosome occupancy across 24,715,362 single molecules.

Individual molecules with the same synthetic reporter sequence display diverse patterns of methylation that correspond to distinct and interpretable molecular binding configurations, including molecules that appear completely covered by nucleosomes (∼147 bp streaks of protection punctuated by ∼50 bp of accessibility), fully or partially bound by TFs (protection at GpCs immediately flanking TetOs), or a mix of the two (**Fig. 1D**). Prior to the induction of TF binding (-dox), the synthetic reporter constructs were relatively inaccessible on average, with methylation probabilities of around 20% (**Fig. 1E**, gray). After 24 hours of dox treatment, we observe a marked increase in accessibility across the reporter constructs and the appearance of clear TF footprints at the positions of TetO sites (**Fig. 1E**, black).

To assign molecular states to single molecules, we developed a probabilistic model that identifies the most likely configuration of TF binding and nucleosome positions given the methylation signal (**Fig. 1D, Fig. S1H, Fig. S2** and Methods). After applying this model to all molecules in our dataset, we observed substantial heterogeneity in the molecular configurations even on identical enhancer constructs, with variable nucleosome positions and degrees of TF binding (**Fig. 1F**, **Fig. S1I-J**). This heterogeneity in TF binding is expected from statistical mechanics and cannot be explained by a fast TF off-rate combined with irreversible methylation that would decrease observed binding (**Fig. S1K**). We also observe substantial variation in nucleosome occupancy over the promoter, identifying two broad classes of promoter states – those with a nucleosome-sized inaccessible region covering the transcription start site (TSS) (which we term “nucleosomal”), and those with nucleosome-free promoters (which we term “active”; **Fig. 1F**, Methods). Within the active promoters, we observe further heterogeneity that likely corresponds to transcriptional machinery (**Fig. S1L**); however, the relative fraction of these states within active promoters does not vary in our assay (**Fig. S1M**). We observe a dox-dependent increase in both the fraction of enhancers that are TF-bound and the fraction of active promoters, although there are many more molecules bound by at least one TF (73% for 5xTetO) than there are molecules with an active promoter (22% for 5xTetO) (**Fig. 1F**).

### TF occupancy exhibits cooperative behavior as a function of the number of binding sites

We set out to determine how the number of TF binding sites affects TF occupancy on individual molecules. Increasing the number of TetO sites resulted in a non-linear increase in the fraction of accessible and TF-bound enhancers, with a large increase between four and six TetO sites (**Fig. 2A**). Furthermore, the average number of bound TFs exhibited cooperative behavior as a function of the number of binding sites (**Fig. 2B**). This average occupancy directly correlates with accessibility as measured by ATAC-seq peak strength (**Fig. S3A**). The TF binding distribution of each enhancer construct, even those with high numbers of sites such as 7xTetO, revealed that very few molecules have all TetO sites occupied (**Fig. 2C**). Instead, we observed a bimodal distribution composed of molecules with no TF binding and a wide distribution of occupancy within TF-bound molecules (with a mode of 3-4 sites bound for 7xTetO).

**Figure 2:**
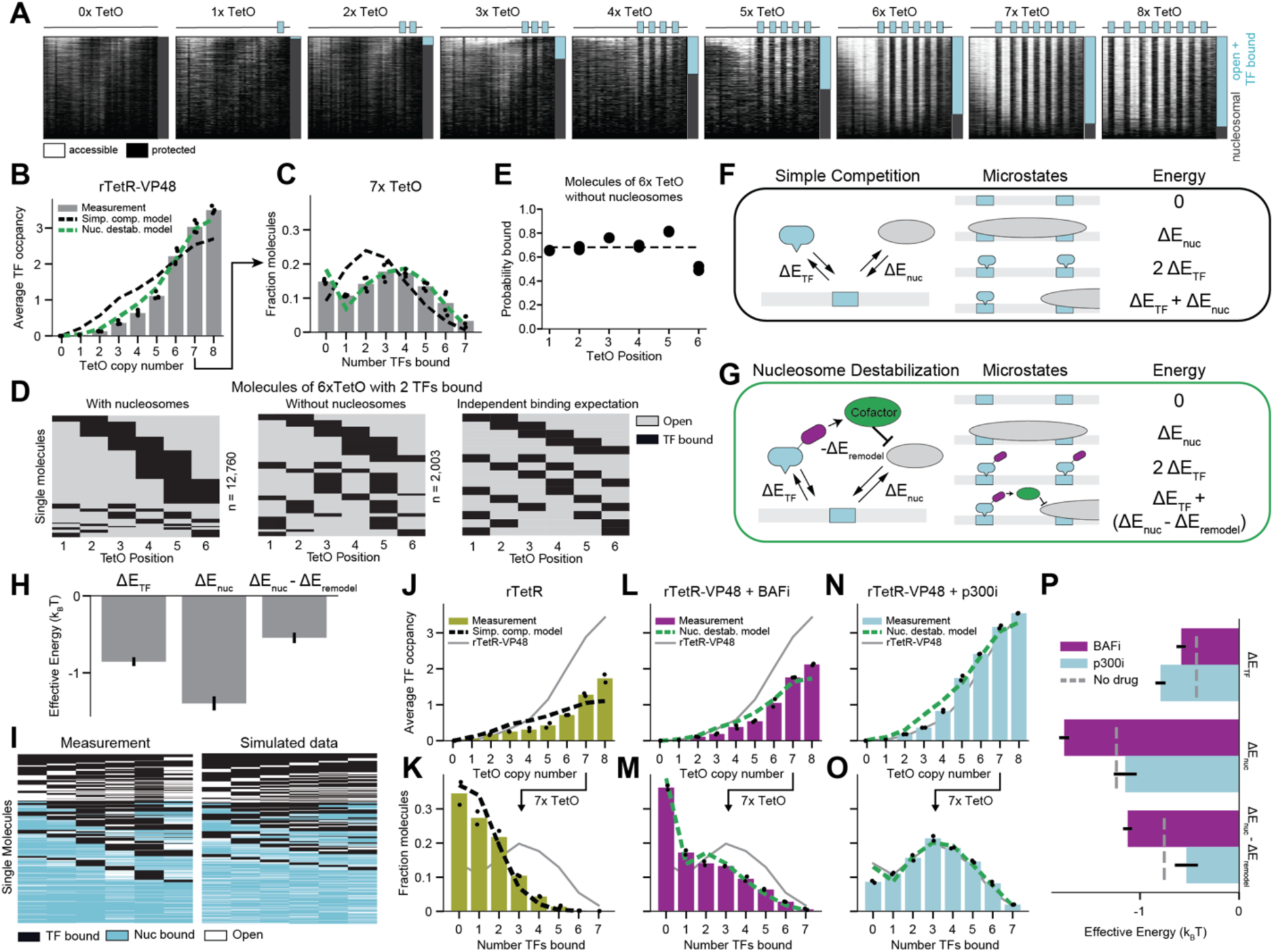
Thermodynamic models reveal rTetR-VP48 competes with nucleosomes with the aid of activation domains. A) Summary plots of enhancer methylation across single molecules observed by SMF for enhancers with 0-8 TetO binding sites after 24 hours of 1,000 ng/ml dox induction. Each row represents methylation data for one molecule, as in 1F. B) Observed average occupancy of rTetR-VP48 for enhancers with increasing numbers of TetO binding sites (four biological replicates) fit to a Simple Competition model (described in F, r^2^ = 0.84) or a Nucleosome Destabilization model (described in G, r^2^ = 0.99). C) Observed occupancy distribution of rTetR-VP48 TFs for the enhancer with 7 TetO sites (four biological replicates) fit to a Simple Competition model (described in F, r^2^ = -0.72) or a Nucleosome Destabilization model (described in G, r^2^ = 0.79). D) Observed distributions of occupied TetO sites on molecules of 6xTetO with 2 TFs bound with and without nucleosomes present and the predictions from a model that assumes independent binding. E) Likelihood of distinct TetO binding sites of being occupied in nucleosome-free molecules containing 6 TetO sites, with position 6 being closest to the promoter (two biological replicates). The dashed line indicates the average occupancy. F) Schematic of the Simple Competition model describing a direct competition of nucleosomes and TFs for binding to DNA. G) Schematic of the Nucleosome Destabilization model describing a TF-dependent recruitment of a cofactor that reduces the binding energy of a nucleosome when one or more TFs are present. H) Best fit parameters and error (SEM) for binding of rTetR-VP48 using Nucleosome Destabilization Model in (G). I) Full molecular state representations for 10,000 measured (left) or simulated (according to the Nucleosome Destabilization, right) 6xTetO enhancer molecules, where each column represents a TetO site and each row represents a molecule. Sites are colored by their occupancy status. J-K) Observed average occupancy (J) and occupancy distribution (K) of rTetR (only) (two biological replicates) with a fit from the Simple Competition model (r^2^ = 0.77 and 0.94, respectively). Gray line is rTetR-VP48 data for comparison. L-M) Observed average occupancy (L) and occupancy distribution (M) of rTetR-VP48 in the presence of the BAF inhibitor BRM014 (two biological replicates) with a fit from the Nucleosome Destabilization model (r^2^ = 0.94 and 0.96, respectively). Gray line is rTetR-VP48 data for comparison. N-O) Observed average occupancy (N) and occupancy distribution (O) of rTetR-VP48 in the presence of the p300 inhibitor A485 (two biological replicates) with a fit from the Nucleosome Destabilization model (r^2^ = 0.99 and 0.90, respectively). Gray line is rTetR-VP48 data without inhibitors for comparison. P) Parameter fits for models in L-O. Error bars represent the standard error of the mean from two biological replicates. Gray dashed line is rTetR-VP48 parameter fits for comparison.

We then investigated the arrangement of individual TF binding events on the same molecule (i.e. where two TFs are bound over six possible sites). We see a strong preference for TFs bound to adjacent sites on molecules with nucleosomes, which represent the majority of molecules in our dataset (**Fig. 2D**). However, once we condition on molecules with no nucleosomes over the enhancer, we observe equal representation of molecular configurations (**Fig. 2D**) that is consistent with independent binding on free DNA. This clustering is expected based on statistical mechanics, without having to invoke direct TF-TF interactions: since nucleosomes span multiple TetO sites, states with adjacent TF binding allow more nucleosomes to bind to that DNA, leading to a more favorable energy state and thus a higher probability of being observed. Furthermore, we do not observe positional biases along the molecule and find that TetO sites are largely (but not always) bound on free DNA (**Fig. 2E**).

### TF occupancy is inconsistent with simple nucleosome competition

We tested if these observed molecular configurations at enhancers can be described with a thermodynamic partition function model, which takes as input effective binding energies associated with protein-DNA interactions of TFs and nucleosomes and predicts the distribution of binding microstates in cells. For each enhancer construct, we identified the set of binding microstates by enumerating all possible configurations of nucleosomes and TFs on DNA and computed their Boltzmann probabilities by summing the effective energies of each molecular interaction (Methods)^18–20^. This model assumes that TF binding is independent and is at (non-equilibrium) steady state, which we confirmed by showing that rTetR-VP48 binding distributions do not change between 24 and 48 hours (**Fig. S3B**). We first tested the simplest thermodynamic model where TFs and nucleosomes compete for the same DNA which we name “Simple Competition Model”. This two-parameter model, which allows only for TF-DNA and nucleosome-DNA interaction energies (**Fig. 2F**), cannot reproduce the bimodality observed in our binding data (**Fig. 2C**).

### TF-dependent recruitment of nucleosome-destabilizing cofactors explains observed molecular states

Because VP48 is known to directly interact with chromatin-remodeling cofactors^21–23^, we modified the model to include a term representing a remodeler-dependent decrease in nucleosome affinity upon TF binding (which we name “Nucleosome Destabilization model”) (**Fig. 2G**). This three parameter model accurately recapitulates the fraction of TF-bound molecules, the average TF occupancy per molecule, and the TF binding distributions across enhancer constructs (**Fig. 2B-C,H, Fig. S3C-D**). Alternative models encoding other VP48-dependent molecular interactions are unable to similarly fit the observed data (**Fig. S3E**). We used the Nucleosome Destabilization model to generate predicted distributions of binding microstates which were qualitatively similar to those observed in our measurements (**Fig. 2I**).

To test the assumptions of our model, we first carried out the same SMF measurements in a cell line expressing rTetR without VP48 (Methods), thus unable to recruit coactivators, and observed an average reduction in TetO occupancy of 35% across the library (**Fig. 2J-K)** that could not be explained by a reduced TF concentration (**Fig. S3F**). The Simple Competition model (without the nucleosome destabilization term) fits these data well (**Fig. 2J-K**, **Fig. S3G-J**), suggesting that mass action kinetics alone is sufficient to explain binding competition between nucleosomes and DNA binding domains that are unable to recruit cofactors. This also implies that the binding of TFs is not simply determined by their DNA-binding domains, but rather that activation domains also play a large role in determining *in vivo* TF occupancy.

### BAF, but not p300, contributes to nucleosome destabilization at the enhancer

To determine the molecular mechanisms underlying the observed nucleosome destabilization upon TF binding, we repeated these experiments in the presence of inhibitors of two major coactivators, starting with the chromatin remodeler BAF. We used the small molecule BRM014, a catalytic inhibitor of the ATPase activity of BRG1/BRM^24^. BAF inhibition reduced the binding of rTetR-VP48, decreasing the fraction of accessible enhancers (**Fig. S3I**), the average TF occupancy per enhancer (**Fig. 2L**), and the TF binding distribution per molecule (**Fig. 2M**) compared to rTetR-VP48 without inhibitors (**Fig. 2B-C, Fig. S3C**). BAF inhibition did not affect the binding of rTetR alone (**Fig. S3K**), suggesting that BAF is specifically recruited by VP48^21^. When we fit the Nucleosome Destabilization model to the BAF inhibitor data, we found that the TF-dependent nucleosome destabilization energy and the nucleosome energy were both decreased, consistent with expectations that BAF activity would influence apparent nucleosome affinity (**Fig. 2P**). The 2 parameter Simple Competition model does not completely fit these data (**Fig. S3L**), suggesting incomplete BAF inhibition, redundant chromatin remodelers that can compensate for loss of BAF^25^, or that activation domains augment TF binding through additional mechanisms independent of chromatin remodelers.

We next tested if the catalytic activity of p300, another enzymatic cofactor that interacts with VP48^26,27^, might also be responsible for nucleosome destabilization upon TF binding. p300 occupancy and p300-catalyzed acetylation of histone H3 lysine K27 are commonly observed at native enhancers^28^, and it has been shown that acetylated histones have weaker affinity to DNA *in vitro*^29^. We inhibited p300 catalytic activity with the small molecule A485^30^, but did not observe a decrease in TF binding (**Fig. 2N-O** vs. **Fig. 2B-C**) or nucleosome occupancy (**Fig. S3M**), despite a marked reduction in H3K27 acetylation (**Fig. S3N**). Moreover, using the Nucleosome Destabilization model to fit these data shows that the nucleosome remodeling energy is not decreased by the p300 inhibitor compared to no inhibitor (**Fig. 2P**), suggesting that p300 catalytic activity is not responsible for TF-mediated nucleosome destabilization in this system.

### Multiple bound rTetR-VP48s contribute additively to promoter activation in a time-averaged manner

We next examined the relationship between promoter states and variable numbers of TF binding sites. Similar to enhancer accessibility, increasing the number of TetO sites results in a non-linear increase in promoter accessibility and the fraction of promoters classified as active (nucleosome-free) upon 24 hours of dox induction (**Fig. 3A**). To control for possible sequence context effects on gene expression, we repeated the experiment with two additional TetO libraries (referenced as backgrounds 1 and 2) with distinct sequences between binding sites from the library used thus far (background 0). In addition, we aimed to saturate gene expression by including up to 9 TetO sites (**Fig. S4A**), necessitating a slight decrease in TetO spacing (19 vs. 21 bp). Across all three backgrounds, we observe non-linear increases in promoter activation (**Fig. 3B**) and TF binding across TetO copy numbers (**Fig. 3C, Fig. S5A-C**), although the magnitude of both is variable with sequence context. However, the fraction of active promoters is always proportional to the average number of bound TFs with an invariable slope across all backgrounds (**Fig. 3D**), demonstrating that each additional bound TF contributes the same amount to promoter activation. Removing VP48 eliminates active promoters (**Fig. S4B**) but does not completely ablate TF binding (**Fig. 2J-K**), indicating that the active promoter fraction is not simply a consequence of binding-induced nucleosome repositioning. Furthermore, inhibiting transcription with either α-amanitin or flavopiridol results in a lower fraction of active promoters, but does not influence the average occupancy of rTetR-VP48 (**Fig. S4C-D**), demonstrating that transcription does not feed back on rTetR-VP48 occupancy.

**Figure 3:**
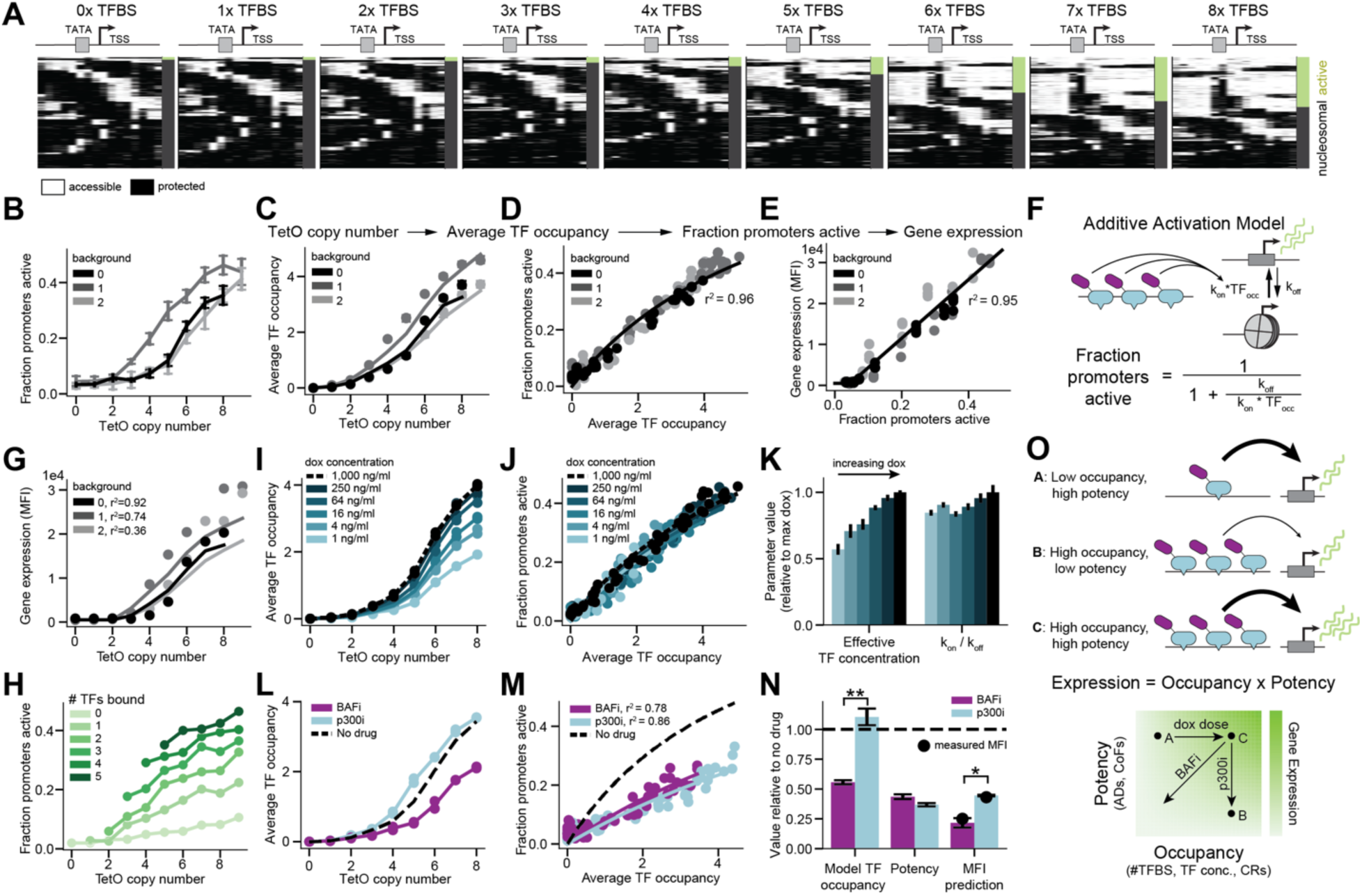
Average rTetR-VP48 occupancy predicts nucleosome free promoters and gene expression. A) Summary plots of promoter methylation of single molecules for enhancers with 0-8 TetO binding sites after 24 hours of dox induction. B) Fraction active (nucleosome-free) promoters as a function of the number of TetO sites across three DNA background sequences. C) Average rTetR-VP48 occupancy as a function of TetO binding sites for three DNA backgrounds. Lines represent the predictions from the Nucleosome Destabilization model fit on each background separately. D) Relationship between rTetR-VP48 occupancy and the fraction of active promoters across three backgrounds. Black line is an additive activation model fit from E (kon/koff = 0.15 ± 0.002 TF^-1^). E) Relationship between fraction of active promoters and gene expression by flow cytometry for three DNA backgrounds. Black line is a thresholded linear fit. F) Schematic of an additive kinetic model whereby TFs contribute independently to promoter activation (kon) by modulating the rate of transitioning from the promoter off state to the on state (Additive Activation model). G) Relationship between TetO copy number and gene expression by flow cytometry for three DNA backgrounds. Lines are combined model fits (Nucleosome Destabilization model, Additive Activation model, and promoter to gene expression model) inputting TetO copy number. H) Observed fraction of active promoters for molecules with N (0-5) rTetR-VP48 TFs bound for different numbers of TetO sites. Error bars are bootstrapped standard deviations. I) Relationship between number of TetO binding sites and the average rTetR-VP48 occupancy and for different concentrations of dox (after 24 hours of exposure). J) Relationship between average rTetR-VP48 occupancy and the fraction of active promoters for different concentrations of dox (after 24 hours of exposure). Lines are Additive Activation model fits. All r^2^ between 0.84-0.97. K) Parameter values of effective rTetR-VP48 concentration (Nucleosome Destabilization model) and potency (Additive Activation model) across dox concentrations relative to maximum dox. L) Relationship between number of TetO binding sites and the average rTetR-VP48 occupancy under conditions of BAF inhibition (purple) or p300 inhibition (cyan). Dotted line is data without inhibitors. M) Relationship between average rTetR-VP48 occupancy and the fraction of active promoters under conditions of BAF inhibition (purple) or p300 inhibition (cyan). Lines are Additive Activation model fits. Dotted line is best fit to data without inhibitors. N) Quantification of average TF occupancy from Nucleosome Destabilization model, potency from Additive Activation model, and prediction of gene expression by coupling these models under conditions of BAF inhibition (purple) or p300 inhibition (cyan) relative to fits without inhibitors for molecules with 5-8 TetO sites in background 0. Black dots for gene expression are measured MFI from flow cytometry on individual cell lines. Significance is determined by paired T-test with 4 degrees of freedom (t-valueoccupancy=-15.8, t-valueMFI=-7.4). O) Schematic showing that parameters of TF occupancy (including #TFBS, TF concentration, and chromatin remodelers (CRs)) and TF potency (including activation domains (ADs) and cofactors (CoFs)) separably tune gene expression.

We then observed a linear relationship between the fraction of active promoters and gene expression, both by flow cytometry for Citrine fluorescence (**Fig. 3E**) and RT-qPCR for the Citrine transcript **(Fig. S4E**), demonstrating that the fraction of active promoters can be interpreted as an accurate single-molecule metric for transcriptional output. This relationship between TF occupancy and gene expression can be well fit by modifying the commonly-used “telegraph” model of transcription^31–33^, which assumes that promoters can be either “on” (transcribing) or “off”, to include a promoter on-rate that is proportional to the average number of TFs bound (Additive Activation model; **Fig. 3F**, Methods). That each bound TF contributes additively to the rate of promoter activation suggests that, for this system, the observed non-linearity between gene expression and TetO copy number stems primarily from AD-dependent nucleosome destabilization via chromatin remodelers (allowing for increased TF binding) and not from additional synergy in promoter activation once TFs are bound (**Fig. S4F-H**). In combining our models for TF occupancy from TetO number (Nucleosome Destabilization model, 3 parameters for each background), promoter activation from TF occupancy (Additive Activation model, 1 parameter across all backgrounds), and gene expression from promoter activation (linear fit, 1 parameter across all backgrounds), we can largely explain gene expression as a function of the number of TetO sites (**Fig. 3G**).

Given that average TF occupancy predicts average promoter activity remarkably well, we next asked if the instantaneous occupancy state at the enhancer can predict the promoter activity state of an individual molecule. The probability that we find a promoter in the active state increases with the number of TFs bound on the same molecule when we combine data across all enhancers (**Fig. S4I)** or restrict to individual enhancers (**Fig. S4J**). However, after conditioning on molecules with the same number of TFs bound, there is still a dependence on the number of TetO sites (**Fig. 3H**). The finding that promoter state is not solely determined by instantaneous TF occupancy is consistent with a model in which the timescales of TF binding and promoter activation are similar (compare **Fig. 3H** to **Fig. S4K** for dissimilar timescales) and suggests that promoters integrate information from multiple rounds of TF binding^34,35^.

### rTetR-VP48 binding and potency can be independently modulated to tune promoter activation

The contribution of each bound TF to increasing promoter on-rates (the slope in **Fig. 3D**) might be considered a steady-state “potency”: a TF’s capacity to activate transcription once bound. To test if this relationship between TF occupancy and promoter activation holds independent of perturbations to binding, we performed SMF again under different dox concentrations. By reducing the dose of dox, we tuned the effective rTetR-VP48 concentration^36^ as measured *in vitro* by an Electrophoretic Mobility Shift Assay (**Fig. S4L-O**), and observed a dox-dependent decrease in the average TF occupancy across TetO copy number (**Fig. 3I, Fig. S4P**). However, average TF occupancy continues to linearly contribute to the promoter activation rate across dox concentrations with the same slope (**Fig. 3J**). From the parameters fit from our Nucleosome Destabilization and Additive Activation models, we observe that reducing dox concentration leads to monotonically decreasing apparent TF concentration (Methods) and little change to the promoter on-rates (**Fig. 3K**). This demonstrates that potency is an intrinsic property of a TF that is independent of a TF’s ability to bind DNA.

BAF inhibition decreases average TF occupancy while p300 inhibition has little to no effect (**Fig. 3L**). In contrast, both BAF and p300 inhibition decrease TF potency as fit by the Additive Activation model (**Fig. 3M**), demonstrating that this potency depends on the activity of the cofactors that VP48 recruits. As effects on occupancy and potency both impact expression, our combined TF binding and promoter activation models correctly predict that BAF inhibition has a larger effect on expression than p300 inhibition (**Fig. 3N**). Overall, these results demonstrate that average TF binding occupancy and TF potency are separable parameters that multiplicatively predict transcriptional output (**Fig. 3O**).

### Single-molecule footprinting can report distinct occupancy states at interferon-stimulated response elements

We next investigated how the general principles connecting TF occupancy and potency to promoter state apply when gene activation is induced by the binding of the endogenous trimeric TF complex ISGF3 (IRF9:STAT1:STAT2) central to mounting an immunological defense to viral infection. ISGF3 mirrors aspects of the rTetR-VP48 system: constitutively expressed subunits rapidly activate gene expression after induction by type I interferons (IFNs), and its motifs are frequently clustered within 500 bp of IFN-stimulated gene (ISG) TSSs (**Fig. S6A**). Previous investigation of this pathway suggested that upon binding of type I IFNs to cell surface receptors, activated JAK-STAT kinases phosphorylate STAT1/STAT2 and enable the ISGF3 trimer to form, enter the nucleus, and bind IFN-stimulated response elements (ISREs)^37,38^. However, recent studies based on ChIP data^39^ propose a model for ISGF3 assembly wherein IRF9 (with possible co-binding of STAT2) is bound to ISREs pre-stimulation and upon stimulation, phosphorylated STAT1:STAT2 assembles with IRF9 on DNA to form the transcriptionally active trimer (**Fig. 4A**).

**Figure 4:**
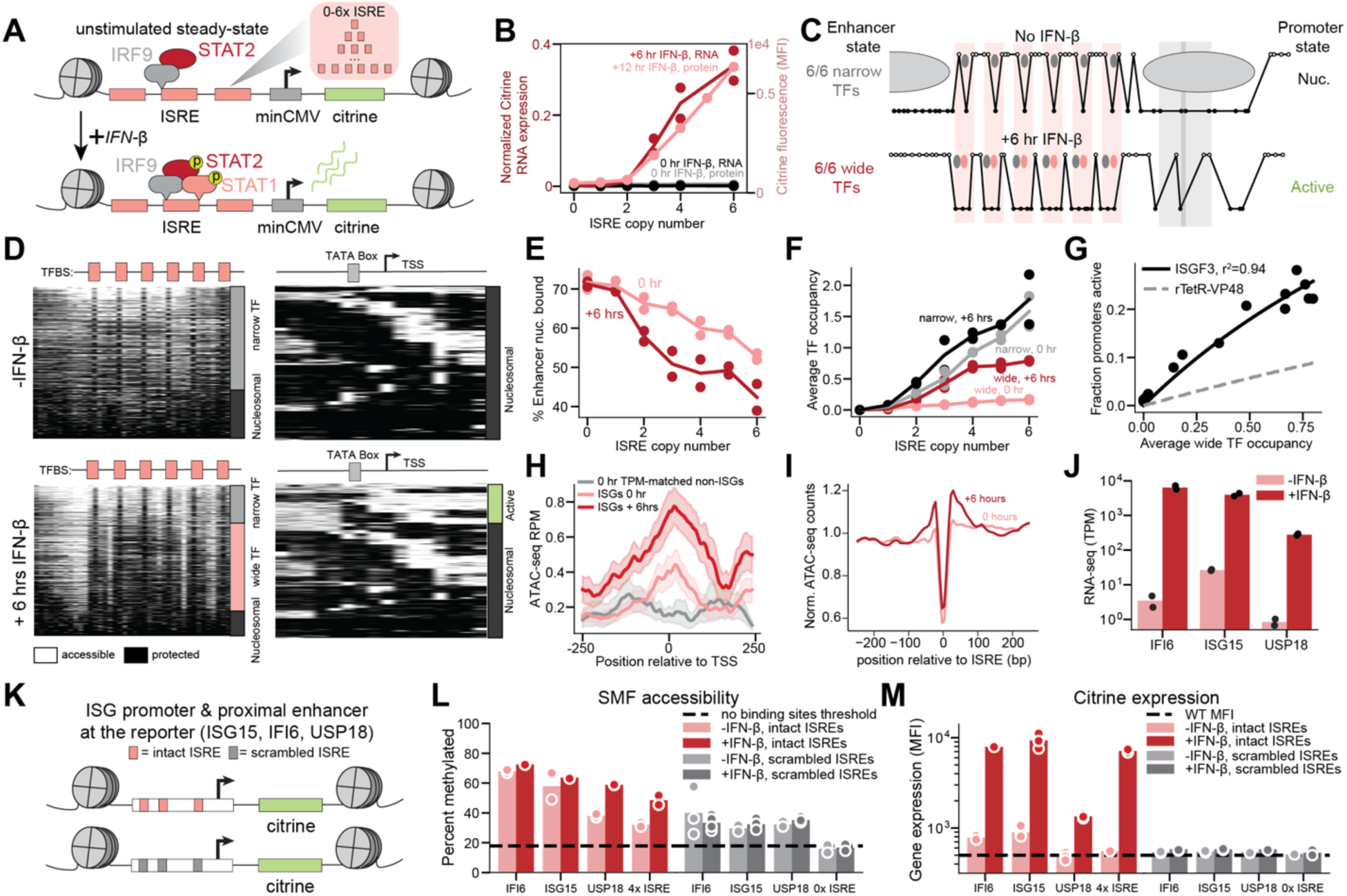
Single-molecule footprinting of an type I interferon response reporter reveals decoupling of accessibility and activation. A) Variable numbers (0-6) of interferon-stimulated response elements (ISRE, pink) upstream of a minCMV (gray) promoter and Citrine gene (green) were integrated at the AAVS1 locus in K562 cells. In the unstimulated state, we expect a IRF9 monomer or IRF9-STAT2 heterodimer to bind the ISRE. Upon IFN-ꞵ stimulation, we anticipate binding of the IRF9-phosphoSTAT1-2 trimer and expression activation. B) Gene expression (as measured by RT-qPCR of Citrine mRNA) and protein levels (as measured by flow cytometry of Citrine) as a function of the number of ISREs before and after IFN-ꞵ stimulation (6 hours of stimulation for RNA and 12 hours for protein) for two technical replicates. C) Exemplar SMF data displaying six narrow footprints in the absence of stimulation (top), and six wide footprints in the presence of IFN-ꞵ (bottom). D) Summary plots of single molecules observed by SMF for enhancers (left) and promoters (right) of constructs with six ISREs without (top) and with (bottom) 6 hours of IFN-ꞵ induction. Each GpC spans an equal width in this representation. The fraction of enhancer sites with majority narrow TF footprints (gray), wide TF footprints (pink) and fully nucleosomal (black) is shown, as is the fraction of promoters that are not bound by a nucleosome (active; green) vs bound by a nucleosome (black). E-F) Observed occupancies of nucleosomes (E) and wide and narrow TF footprints (F) as a function of the number of ISREs before and after six hours of stimulation for two biological replicates. G) Relationship between wide footprint occupancy and the fraction of active promoters for two biological replicates. Black line is an Additive Activation model fit (see 3G, kon/koff = 0.44 ± 0.02 TF^-1^). Gray dotted line represents the model fit for rTetR-VP48. H) Average ATAC-seq accessibility over two biological replicates before (pink) and after six hours (red) of stimulation relative to the TSS of genes upregulated with stimulation (as identified by bulk RNA-seq) that contain at least three ISREs within the 500 bp window. Gray line is average ATAC-seq accessibility for non-ISG promoters that are matched to the pre-stimulation expression level (TPM) of plotted ISG promoters. Shaded error regions are 95% confidence intervals from bootstrapping. I) Average ATAC-seq accessibility over two biological replicates relative to ISREs genome-wide, Tn5-bias corrected and RPM normalized. J) RNA expression of endogenous ISGs before (pink) and after (red) stimulation over two biological replicates of RNA-seq. K) Promoters and promoter-proximal enhancers of ISGs IFI6, ISG15 and USP18 containing ISREs (pink) as well as perturbations scrambling only the ISREs (gray) upstream of Citrine gene (green) were engineered into the AAVS1 locus in K562 cells. L) Average SMF accessibility of ISG promoter reporters with intact ISREs pre-stimulation (pink) and after six hours of stimulation (red) and scrambled ISREs pre-(light gray) and post-stimulation (dark gray) for two biological replicates. Two versions of scrambled ISREs were measured for each construct. Black dashed line is the average SMF accessibility of a sequence with no intentional binding sites. M) Gene expression (as measured by flow cytometry) of ISG promoter reporters with intact ISREs pre-stimulation (pink) and after 24 hours of stimulation (red) and scrambled ISREs pre-(light gray) and post-stimulation (dark gray) for two technical replicates. Two versions of scrambled ISREs were measured for each construct. Black dashed line is the MFI of WT K562s.

To examine the molecular events underlying type I IFN response, we installed a library of promoter-proximal enhancers containing variable ISRE copy numbers upstream of minCMV and the Citrine reporter in K562s (**Fig. 4A**). We observe a non-linear increase in reporter expression (Citrine fluorescence and mRNA levels) with increasing ISRE copy number only after addition of IFN-ꞵ (**Fig. 4B**). After confirming that the kinetics of ISGF3 binding were suitable for footprinting (**Fig. S6B**), we generated SMF data for conditions without IFN-ꞵ and after six hours of IFN-ꞵ stimulation. Despite the lack of reporter expression, TF footprints and nucleosome-free regions are present at ISREs pre-stimulation, consistent with the model of pre-bound but transcriptionally inactive IRF9^39^ (**Fig. 4C-D, Fig. S6C**). Upon stimulation, footprints at ISREs become wider (**Fig. 4C-D**), accessibility increases (**Fig. 4D**), and fewer nucleosomes are bound across the enhancer and promoter (**Fig. 4E**). Together, these findings imply that the wider footprints (due to increased steric blockage of methylation at ISREs) correspond to ISGF3 trimer binding and are consistent with the known structure and DNA contacts of ISGF3 and its subunits^40,41^. Given this ability to resolve binding patterns of distinct molecular configurations, we updated our binding model to distinguish between narrow (IRF9) and wide footprints (ISGF3) at ISREs (**Fig. 4C, Fig. S7**). TF occupancy increases with the number of binding sites for both the narrow and wide footprints, although we observe appreciable wide footprints only upon stimulation (**Fig. 4F**). Before stimulation, the TF binding distribution of narrow footprints is better fit by the Nucleosome Destabilization than the Simple Competition model (**Fig. S6D-G**), suggesting that IRF9 alone may interact with chromatin remodelers to decrease local nucleosome affinity. Furthermore, induced reporter expression is ablated by BAF inhibition (**Fig. S6H**), consistent with previous findings that chromatin remodelers are a key component in type I IFN response for both basal and induced expression^42^.

### TF footprints explain ISRE-driven basal and stimulated accessibility and expression

Similar to our findings with rTetR-VP48, the IFN-ꞵ-stimulated fraction of active promoters increases non-linearly with increasing ISRE copy number (**Fig. S6I**). The upregulated active promoter states are similar to those in the fully synthetic system (**Fig. S6J**) and we do not observe them in the absence of IFN-ꞵ (**Fig. 4D**), consistent with the observation that rapid induction in IFN response is not due to the presence of paused polymerase^43^. While narrow footprints are not predictive of the fraction of promoters active **(Fig. S6K)**, the average number of wide footprints (active ISGF3) predicts the fraction of active promoters, is well fit by the Additive Activation model, and demonstrates a stronger potency (slope) than rTetR-VP48 (**Fig. 4G**). Furthermore, the fraction of active promoters remains linearly related to gene expression (**Fig. S6L)**, demonstrating that expression level remains directly proportional to the aggregate occupancy of activating TFs (**Fig. 4SM**).

Given these results, we hypothesized that ISREs at endogenous ISG regulatory elements are responsible for driving both pre-stimulation chromatin accessibility and post-stimulation transcription. Genome-wide measurements of accessibility (ATAC-seq) and expression (RNA-seq) demonstrate that IFN-ꞵ initiates the expected response in K562 cells, with motif enrichments in differentially accessible sites for IFN response factors and upregulation of known ISGs (**Fig. S6N-O**). Consistent with our hypothesis, the promoters and promoter-proximal enhancers of ISRE-containing ISGs demonstrate more pre-stimulation accessibility in ATAC-seq compared to non-ISGs (**Fig. 4H**), and we observe pre-stimulation TF footprints at ISREs genome-wide (**Fig. 4I**). To determine the degree to which ISREs regulate both accessibility and expression of endogenous ISGs, we decided to further investigate three significantly upregulated genes: ISG15, IFI6, and USP18 (**Fig. 4J**). We installed the core promoter and promoter-proximal ISRE-containing enhancers of these ISGs upstream of the Citrine reporter (**Fig. 4K**) and performed SMF. These regulatory elements exhibit significant pre-stimulation accessibility with variable increases post-stimulation (**Fig. 4L, Fig. S6P**), consistent with ATAC-seq data at the endogenous genes (**Fig. S6Q**), and only express Citrine post-stimulation (**Fig. 4M**). When we scramble only the ISREs, both the pre-existing accessibility and stimulated gene expression are ablated (**Fig. 4L-M**), demonstrating that ISREs are the critical sequence elements for driving and decoupling accessibility and expression at ISGs.

### Temporal delay between rTetR-VP48 binding and promoter activation confirms promoters integrate over multiple binding events

The rapid transcriptional activation of ISGs has likely emerged from the evolutionary pressure exhibited on Type I IFNs^44,45^ to quickly and efficiently fight viral infection. Because we observed both higher pre-stimulation TF binding and a stronger potency for ISGF3 compared to rTetR-VP48, we wanted to investigate what determines activation dynamics across both systems. Assuming that the promoter off-rate is intrinsic to the minCMV promoter (used with both TetO and ISRE enhancer libraries), our measured TF potency (kon/koff) makes strong predictions about activation dynamics. Since steady-state TF binding and DNA accessibility are higher pre-activation in the interferon system compared to the rTetR-VP48 system, TF binding dynamics are also likely to differ.

To test these hypotheses, we stimulated rTetR-VP48/TetO reporter lines with saturating dox for different amounts of time and performed SMF. We observed a time-dependent increase in the binding of rTetR-VP48 which reached steady state by 8 hours, although a significant portion of the TF binding occurs while cells are being processed in buffers containing dox during the assay (**Fig. 5A, Fig. S8A**). However, we observed an hours-long delay between rTetR-VP48 binding and transcriptional activation (as measured by both the active promoter fraction and RT-qPCR), which does not reach steady state until between 8 and 24 hours (**Fig. 5B, Fig. S8B**). Average rTetR-VP48 occupancy remains linearly related to the active promoter fraction at each timepoint during this activation process (**Fig. 5B**), and the apparent potency increases over time, confirming that promoters integrate information from multiple binding events (**Fig. 5C**).

**Figure 5:**
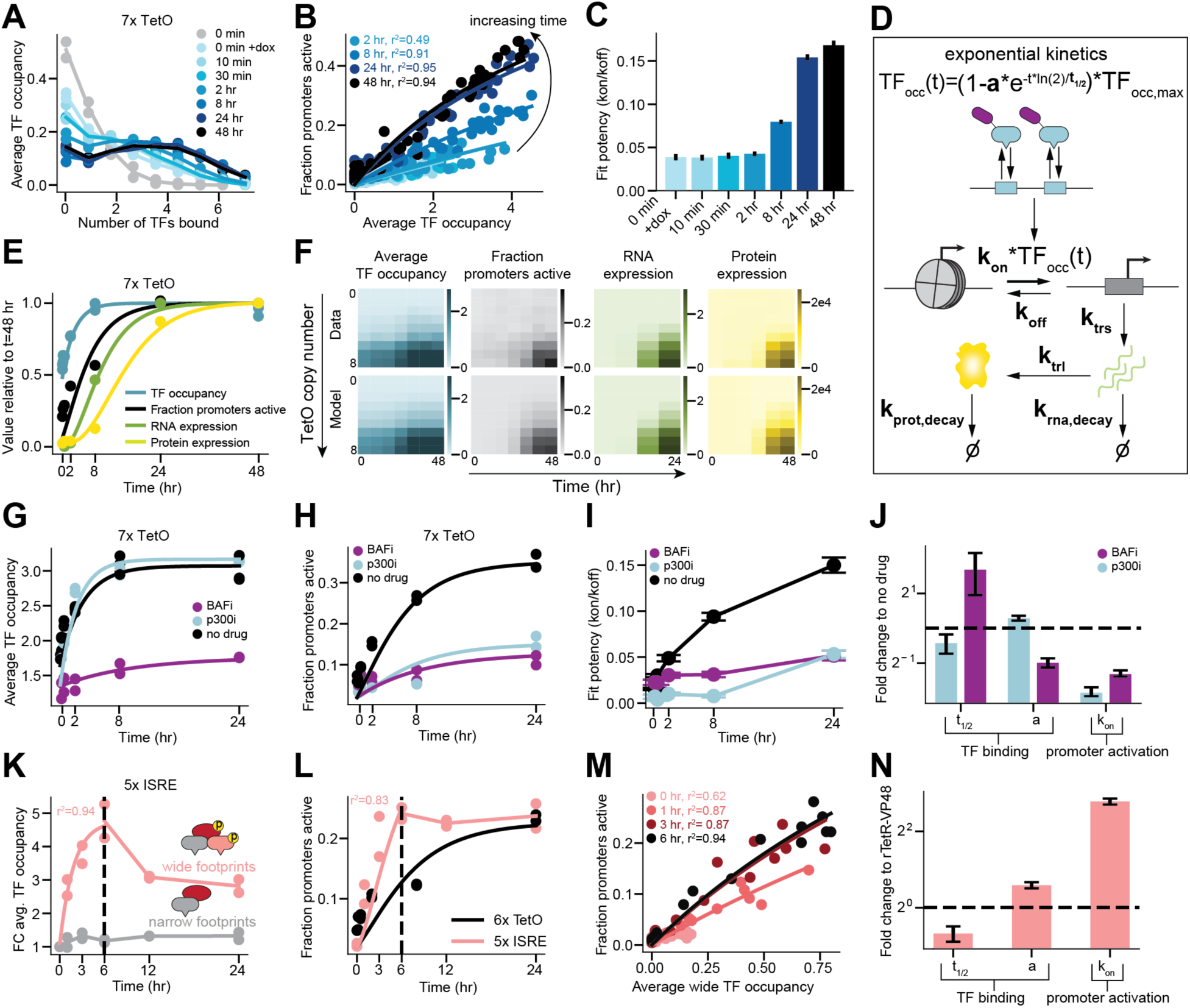
Kinetic modeling reveals timescales for chromatin and gene regulatory changes, as well as relative effects of cofactor inhibition and TF potency on activation. A) The distribution of molecules with different numbers of rTetR-VP48s bound for a construct with 7 TetO binding sites for different times (after dox induction) and two biological replicates. The 0 minute sample (gray) is processed without dox present in buffers during the SMF assay (see Methods) whereas the 0 minute + dox (light blue) sample and all later timepoints are processed with dox present in buffers. B) The relationship between average number of TetO sites bound by rTetR-VP48 and the fraction of promoters active for different times after dox induction and two biological replicates. Lines are Additive Activation model fits. C) Potency (kon/koff) fit from Additive Activation Model across dox timecourse. Error bars are standard deviations. D) A kinetic model for enhancer equilibration, promoter activation, transcription and translation. E) Fits (lines, using kinetic scheme in D) on data (points) for average occupancy of rTetR-VP48, fraction active promoters, RNA expression, and MFI for a construct with 7 TetO binding sites over time. TF occupancy parameters: t1/2 = 2.1 ± 0.3 hr and a = 0.52 ± 0.02. Promoter activation parameters: kon = 0.11 ± 0.02 TF^-1^ hr^-1^ and koff = 0.16 ± 0.04 hr-^1^. RNA parameters: ktrs = 2.0 ± 0.4 RNA (A.U.) promon^-1^ hr^-1^ and kdecay,rna = 0.20 ± 0.05 hr^-1^. Protein parameters: ktrl = 1,200 ± 200 protein (A.U.) RNA (A.U.)^-1^ hr^-1^ and kdecay,prot = 0.16 ± 0.04 hr^-1^. F) Measured data (top) and fit kinetic models (bottom, using kinetic scheme in D) for TF occupancy (r^2^ = 0.99), promoter activation (r^2^ = 0.90), RNA (r^2^ = 0.95), and protein levels (r^2^ = 0.92) over time and across TetO copy number. G-H) Data and fits (using kinetic scheme in D) for TF occupancy (G) and fraction of active promoters (H) for the 7xTetO constructs over time under p300 inhibition (cyan) and BAF inhibition (purple) for two biological replicates. I) Potency (kon/koff) fit from Additive Activation Model across dox timecourse with p300 (cyan) and BAF inhibition (purple). Error bars are standard deviations. J) Kinetic parameters (using kinetic scheme in D) for data in G and H displayed as fold change compared to no drug conditions. Error bars are standard error. K) Fold change of wide footprints (red) and narrow footprints (gray) for a construct with 5x ISRE elements as a function of time after stimulation with model fit up to six hours for two biological replicates. L) Fraction of promoters active for a 5x ISRE enhancer (pink) plotted as a function of time after stimulation with model fit up to six hours for two biological replicates. For comparison to a TetO promoter with the same maximum fraction active, the fraction of promoters active for a 6xTetO enhancer (black) is plotted as a function of time after maximum dox stimulation with model fit for two biological replicates. M) The relationship between average number of ISREs exhibiting a wide footprint and the fraction of promoters for different times after stimulation (up to 6 hours) and two biological replicates. N) Kinetic parameters (using kinetic scheme in D) for data in K and L displayed as fold change compared to parameters obtained for rTetR-VP48. Error bars are standard error.

### Kinetic modeling reveals the molecular basis of kinetic differences between TF activation capacities

Given that average TF occupancy remains predictive of promoter activity over time, we set out to build a model that describes the dynamics of TF binding, promoter activation, transcription, and translation (**Fig. 5D**, Methods). To quantify the rate of rTetR-VP48 binding, we fit a single binding equilibration timescale across enhancers and normalized their TF occupancies to steady-state values (**Fig. 5D-E, Fig. S8C-D**). Extending our Additive Activation model (which assumes that each bound TF increases the activation rate of the promoter), we fit activation dynamics using a set of ordinary differential equations in which the promoter on-rate scales with average TF occupancy (**Fig. 5D-E, Fig. S8E**). Since we observed that the fraction of active promoters is linearly predictive of both mRNA and protein levels at steady state, we used the RT-qPCR and Citrine fluorescence data to fit single rates for transcription, RNA degradation, translation, and protein degradation (**Fig. 5D-E, Fig. S8F-G**). These models accurately capture the temporal delays between successive steps of the central dogma across all enhancers in our rTetR-VP48/TetO reporter library (**Fig. 5E-F**).

Based on our steady-state results with inhibitors (**Fig. 3L-N**), we hypothesized that BAF and p300 inhibition would both reduce the rate of promoter activation, while only BAF inhibition would affect TF binding kinetics. To test this hypothesis, we performed the same timecourse after inhibiting each cofactor. As predicted, BAF inhibition slows down TF binding (**Fig. 5G,J, Fig. S8H**) while both inhibitors decrease the rate of promoter activation (**Fig. 5H,J, Fig. S8I**) and lead to a slower increase in potency (**Fig. 5I**).

Additionally, based on the increased pre-stimulation accessibility and potency of the type 1 IFN reporter, we postulate that ISGF3 would demonstrate both faster TF binding and promoter activation than rTetR-VP48. We performed a similar timecourse on ISGF3 reporter cells. While narrow footprints are constant throughout the timecourse, wide footprints appear rapidly after IFN-ꞵ addition and reach their maximum by six hours before decreasing (**Fig. 5K**), consistent with the known downregulation of type I IFN response after activation^46^. Kinetic model fits (as described in **Fig. 5D**) to wide footprint dynamics reveal a shorter binding half-time than rTetR-VP48, consistent with pre-accessibility enabling faster TF binding (however, a direct comparison cannot be made as this term also depends on TF concentration) (**Fig. 5N, Fig. S8J**). Average wide footprint occupancy remains predictive of the fraction of active promoters during activation, and the promoter dynamics closely follow those of the wide footprints, further suggesting that footprint width can discriminate between inactive IRF9 and active ISGF3 (**Fig. 5L-M**). While promoters remain accessible throughout the timecourse (**Fig. 5L)**, transcription is downregulated (**Fig. S8K)** similarly to the wide footprints (**Fig. 5K)**, in line with the expected downregulation of endogenous ISGs^47^. In contrast to rTetR-VP48, which takes more than eight hours to reach its maximum potency, ISGF3 takes only three, consistent with a shorter delay between TF binding and promoter activation (**Fig. 5B,M, Fig. S8L**). Furthermore, an ISGF3 reporter demonstrates faster activation when compared to an rTetR-VP48 reporter with the same maximum active promoter fraction, and the fit promoter on-rate is >5-fold faster (**Fig. 5L,N, Fig. S8M**). Taken together, the comparison of kinetic parameters between our two systems suggests that IFN-responsive promoters integrate regulatory inputs over less time and that cells have adopted multiple molecular mechanisms to achieve rapid transcriptional activation.

## Discussion

Our study provides high-throughput measurements of protein occupancy at regulatory elements on individual molecules in cells, and explains how TF binding sites, TF identity, TF concentration, and cofactor activity quantitatively tune TF binding and transcription. Our single-molecule occupancy data reveal that regulatory elements exhibit a broad diversity of microscopic binding states, including states that are primarily nucleosome occupied, suggesting that even highly active elements are likely accessible to regulatory proteins only in a subset of cells. However, average single-molecule methylation is linearly correlated with ATAC-seq signal strength, suggesting that genome-wide chromatin accessibility measurements accurately report information regarding the average single-molecule state present in cells. We also identified a decoupling of chromatin accessibility and transcriptional activity in type I IFN response, challenging the common heuristic that accessibility implies transcriptional activity.

The single-molecule binding distributions of both synthetic TFs with activation domains (rTetR-VP48) and native TFs (IRF9) are best fit by a model requiring the destabilization of nucleosome-DNA interactions across the entire regulatory element upon the binding of at least one TF. The necessity of this destabilization parameter, alongside empirical decreases in TF binding upon activation domain removal and cofactor inhibition, implies that activation domain:cofactor interactions play a substantial role in determining TF binding within chromatin. This may help explain the poor prediction of *in vivo* TF occupancy from *in vitro* binding affinities (often using only DNA binding domains)^48^. Furthermore, our quantitative statistical mechanical model also provides a clearly defined mechanism by which a TF might begin to establish accessibility at a regulatory element and facilitate the binding of additional TFs, even without directly binding to nucleosomal DNA.

Our observation that p300 catalytic activity does not contribute to destabilization of nucleosome-DNA interactions appears inconsistent with a model by which histone acetylation reduces nucleosome-DNA affinity to facilitate TF binding^29,49,50^ and is instead consistent with recent observations that inhibiting p300 catalytic activity actually stabilizes interactions with chromatin^51,52^. The observation that inhibiting p300 does have strong effects on promoter activation also supports previous reports on the role of the catalytic activity of p300 in assembling the pre-initiation complex (PIC) and recruiting RNA polymerase^51^. In contrast, we observe that BAF influences both the capacity of TFs to bind (by increasing nucleosome eviction) and acts to facilitate promoter activation (perhaps through evicting nucleosomes that block the TSS). Intriguingly, the distinct effects on binding and promoter activation upon BAF and p300 inhibition are mirrored in their differential effects on the shapes of the flow cytometry distributions (i.e. fraction of ON cells and MFI of ON cells) (**Fig. S4Q**), a phenomenon which deserves further investigation.

We observe that each bound TF adds linearly to the rate of promoter activation^2^, and cooperativity between activation domains in controlling the transcription cycle is not required to explain the synergy between motif copy number and transcription^9^. Furthermore, the instantaneous binding of TFs does not predict the promoter state, but rather promoters integrate information over multiple binding events. Such an integration could occur if promoter on- and off-rates are slower than TF binding, if multiple steps are required to “license” a promoter from an off to on state, or if there exists a molecular integrator of multiple TF binding events (such as histone acetylation)^34,35^.

Comparisons between our synthetic activation system and ISREs suggests that cells have multiple ways to increase the rate by which promoters are activated. One mechanism may be the establishment of pre-accessibility by non-activating TFs (i.e. IRF9), providing a foothold in the chromatin landscape for rapid binding of subsequent activating TFs. These pre-bound factors may also deposit chromatin marks at the promoter that increase its capacity for rapid activation. Alternatively, different TFs may have distinct kinetic capacities for activation, and arbitrating between these hypotheses is an interesting future direction. Uncovering mechanisms to tune the kinetics of transcriptional responses may also enable the design of more complex and robust circuits in synthetic biology.

Our footprinting system also exhibits some salient limitations. First, we only studied two distinct types of TFs that were readily inducible (through dox addition for rTetR-VP48 and through stimulation with IFN-ꞵ for ISGF3). General investigations of TF activities over time are complicated by this requirement for acute induction, as well as the possibility that, in other scenarios, multiple TFs expressed in a single cell type may be binding to similar motifs. Furthermore, the synthetic approach we describe here allowed us to engineer GpC dinucleotides in strategic locations that enabled unambiguous assignment of molecular states across each molecule. The lower abundance and variable positioning of GpCs in the genome makes this task much more challenging at non-engineered genomic loci. Finally, this approach can only be applied to TFs with binding residence times longer than the timescale of the (irreversible) methylation reaction to avoid underestimating the true TF occupancy state at regulatory elements.

Overall, however, our platform for linking microscopic molecular states at a regulatory element to transcriptional states at a promoter provides a facile mechanism to separately quantify the contributions of TF binding and TF potency on gene expression, features that are largely convolved in other assays. Using our dox-inducible system, we anticipate measuring the effects of diverse activation domains fused to rTetR (including potential synergistic interactions between pairs of activation domains) and their cofactor dependencies. Modification of this platform would also enable the investigation of transcriptional repression. Finally, our thermodynamic binding model makes concrete predictions as to the effects of changing TF spacing and binding affinity which we hope to fully explore and validate.

## Methods

### Cell lines and cell culture

Cell culture was performed as described in^53^. Briefly, all experiments were performed in K562 cells (female, ATCC CCL-243). Cells were cultured in a controlled humidified incubator at 37C and 5% CO2, in RPMI 1640 (Gibco, 11-875-119) media supplemented with 10% FBS (Omega Scientific, 20014T), and 1% Penicillin-Streptomycin-Glutamine (Gibco, 10378016). Synthetic transcription factors (rTetR-VP48 and rTetR) were installed into wild-type K562 cells as described in^54^. Briefly, 1 × 10^6^ wild-type K562 cells were electroporated in 100 μl of Amaxa solution (Lonza Nucleofector 2b, program T-016) with 1 µg of PiggyBac expression vector (PB200A-1, SBI) and 1 µg of donor plasmid (pMMH6.2, pMMH107, pMMH090, pMMH096), an ITR-flanked plasmid harboring the EF-1α core promoter driving rTetR-(+/-VP48)-(+/-3xFLAG)-T2A-hygromycin resistance gene and a separate Tet responsive promoter (TRE3G) driving an mCherry gene. All experiments were performed with rTetR(3G) except for FLAG staining which was performed with rTetR(SE-G72P). Integrants were selected to purity using 200 μg ml^−1^ of hygromycin (Thermo Fisher Scientific) over 7 days. Expression level evaluation with FLAG was performed with α-FLAG-Alexa647 (RNDsystems, IC8529R) as in^55^.

Reporter cell lines were then generated as in^55^. Briefly, reporter cell lines were generated by TALEN-mediated homology-directed repair to integrate enhancer reporter donor constructs into the *AAVS1* locus by electroporation of 1 × 10^6^ K562 cells (or K562 cells selected to have the desired synthetic transcription factor) with 1 ng of reporter donor plasmid and 0.5 ng of each TALEN-L (Addgene no. 35431) and TALEN-R (Addgene no. 35432) plasmid. After 2 days, the cells were treated with 1,000 ng ml^−1^ puromycin antibiotic for 7 days to select for a population where the donor was stably integrated in the intended locus. These cell lines were not authenticated. All cell lines tested negative for mycoplasma.

All doxycycline induction was performed by adding 1,000 ng/mL doxycycline (Fisher Scientific, 409050) for 24 hours unless specifically noted otherwise. All interferon induction was performed by adding 10 ng/mL IFN-b (Peprotech, 300-02BC) for 6 hours unless specifically noted otherwise. Brg1 and Brm ATPase inhibition was performed by adding 10 uM BRM014 (MedChem Express, HY-119374) to cells as in^56^ for 30 minutes prior to doxycycline addition. CBP/p300 KAT inhibition was performed by adding 3 uM A-485 (Selleck Chemicals, S8740) as in^57^ for 30 minutes prior to doxycycline addition. Polymerase II inhibition was performed by adding 50 uM alpha-amanitin (Sigma-Aldrich, A2263) for either 6 hours prior to assaying cells or 1 hour prior to doxycycline addition. Transcriptional initiation and elongation inhibition was performed by adding 10 uM flavopiridol (Sigma-Aldrich, F3055) for 1.5 hours prior to assaying cells.

### Flow cytometry and analysis

Flow cytometry data was collected on a BioRad ZE5 flow cytometer and analyzed using Cytoflow^58^ in Python. Live cells were gated by retaining 70% of cells with DensityGateOp. The gate for Citrine ON cells was set as two standard deviations from the mean of a gaussian fit to WT K562 cells (OFF cells).

### RT-qPCR and analysis

We harvested 20,000 cells per condition in duplicate into buffer RLT (Qiagen). We then extracted nucleic acid using Dynabeads MyOne Silane beads (Thermo Fisher), treated samples with TURBO DNase (Thermo Fisher), and cleaned again with the silane beads. We used the SuperScript IV VILOAffinityScript reverse transcriptase master mix (Thermo Fisher) to generate cDNA and performed qPCR using FastStart Universal SYBR Green Master (Sigma Aldrich). We used primers against Citrine (CGGCGACGTAAACGGCCACAAGTTCAG, CTTGCCGGTGGTGCAGATGAA) and GAPDH (AGCACATCGCTCAGACAC, GCCCAATACGACCAAATCC) and calculated effects using the ΔΔCt method. As the expression of metabolic genes is affected during Type I IFN response, the qPCR for the IFN timecourse data are normalized to the GAPDH value prior to stimulation (0 hr).

### Library design

For the rTetR-VP48 reporter libraries (see Supp. Table 1), background 0 was randomly generated with 21 base pairs in between TetO sites, background 1 was a mutated version of background 0 with 19 base pairs in between TetO sites, and background 2 was chosen as a region from hg38 with high GC content that was not in K562 DNase peaks and did not contain simple repeats or key restriction enzyme sites. The maximum number of TetO sites (8 for background 0, 9 for backgrounds 1 and 2) were then inserted with equal spacing and GCs immediately flanking the motif (5’-**gc**TCCCTATCAGTGATAGAGA**gc**-3’). We scrambled the TetOs one at a time from 5’ to 3’ to maintain base pair content. We then ensured that there was at least one GC in between TetO sites and removed all CGs to avoid endogenous methylation. For the ISGF3 reporter libraries (see Supp. Table 1), we generated a library of 0-6x ISREs with 21 base pairs in between motifs in background 0. We placed GCs both immediately flanking and within ISREs (5’-**gc**TAGTTTC**GC**TTTCCC**gc**-3’). All libraries included a control sequence with one CTCF motif inserted (5’-CCACCAGGGGGCGC-3’ - see Supp. Table 1).

### Library cloning

Oligonucleotides of 350 bp in length were synthesized as pooled libraries (IDT), resuspended to 0.1 uM in TE buffer and diluted to 1 nM in H2O. TetO libraries were then PCR amplified to add on SMF primers and Gibson overhangs, while ISRE libraries were ordered with SMF primers and were then PCR amplified to just add Gibson overhangs. For each 50 uL reaction adding SMF primers, we used 1 uL 1 nM library, 2.5 uL each 10 uM forward and reverse primers (cTF254, cTF255) and 25 uL NEBNext Ultra II Q5 Master Mix (NEB, M0544) and ran the following thermocycling protocol: 30 seconds at 98 °C, then 10 cycles of 98 °C for 10 s, 60 °C for 20 s, 72 °C for 40 s, and then a final step of 72 °C for 5 minutes. PCR product was then cleaned with a 1x AmpureXP bead clean and eluted in 15 uL H2O. Gibson overhangs were then PCRed onto the library. For each 50 uL reaction, we used 2.5 uL cleaned product, 2.5 uL each 10 uM forward and reverse primers (cTF256, cTF257) and 25 uL NEBNext Ultra II Q5 Master Mix (NEB, M0544) and ran the following thermocycling protocol: 30 seconds at 98 °C, then 10 cycles of 98 °C for 10 s, 68 °C for 30 s, 72 °C for 20 s, and then a final step of 72 °C for 5 minutes. PCR product was then isolated by running on E-Gel EX 1% Agarose Gels (ThermoFisher G401001), excising the band at the expected length, and extracting with Zymoclean Gel DNA Recovery Kits (Zymo D4007).

Libraries were then cloned into a reporter vector (pCL056) with Gibson assembly. For each 10 uL reaction, we used 25 ng (0.005 pmol) pre-digested and gel-extracted backbone plasmid, 0.01 pmol (2:1 molar ratio of insert:backbone) library and 2.5 uL NEBuilder HiFi DNA Assembly Master Mix (NEB E2621) and incubated at 50 °C for 60 minutes. We next transformed 2 uL of assembled vectors into 10 uL Stable Competent E. coli (NEB, C3040) by placing on ice for 2 minutes, heat shocking at 42 °C for 30 seconds, placing on ice for 2 minutes, and resuspending cells in 950 uL 10-beta/Stable Outgrowth Medium (NEB, B9035S). Serial dilutions of 10 uL were immediately spread onto Ampicillin selection plates to assay transformed library complexity (>= 10,000x). The remainder of transformed cells were incubated shaking at 300 rpm for 30 minutes at 37 °C before they were added to 2 mL of LB with a final concentration of 50 ug/mL Ampicillin (Sigma-Aldrich, A5354) and incubated shaking at 300 rpm overnight. Plasmids were isolated from cultures using the QIAprep Spin Miniprep Kit (Qiagen, 27104).

For creating individual reporter cell lines with minCMV, enhancer reporter donor plasmids were isolated by picking single colonies from above cloning or cloned as above from individual gBlocks (Twist). For creating individual reporter cell lines of ISG regulatory elements and variants, the genomic promoters (up to the TSS) and proximal ISRE-containing enhancer sequences were ordered as gBlocks (Twist, see Supp. Table 1) and cloned as above, with the exception that minCMV of pCL056 was excised in the digest.

### Methylation treatment

Between 1-3 million K562 cells were spun down (5 min, 300 r.c.f., 4°C) in 2mL Eppendorf tubes in a pre-chilled swing-bucket centrifuge, which is used throughout the protocol. The supernatant was aspirated and cells were resuspended in 200uL ice cold PBS. After spinning cells down (5 min, 300 r.c.f., 4°C), the supernatant was aspirated using a P200 to avoid cell loss. To lyse cells using the Omni-ATAC protocol, cells were then resuspended in 200uL of fresh ice cold lysis buffer (10mM Tris-HCl pH 7.5, 10mM NaCl, 3mM MgCl2, 0.1% NP-40, 0.1% Tween-20, 0.01% Digitonin) by pipetting up and down three times. This cell lysis reaction was incubated on ice for 3 minutes. After lysis, 1.2 mL of ice cold resuspension buffer (10mM Tris-HCl pH 7.5, 10mM NaCl, 3mM MgCl2, 0.1% Tween-20) was added, and the tubes were inverted 3x to mix. Nuclei were then immediately spun down (10 min, 500 r.c.f., 4°C). Supernatant was aspirated first with a P1000 and then with a P200 to avoid nuclei loss. Each methylation reaction was thoroughly mixed on ice using 100uL of cell methylation buffer (10% volume 10x M.CviPI reaction buffer (NEB, B0227S), 300 mM sucrose, 2.13 mM S-adenosylmethionine) and 50uL of 4,000U/mL M.CviPI GpC methyltransferase (NEB, M0227L). Each methylation reaction was then incubated for at 37 °C shaking at 1000 rpm for 7.5 minutes (unless specified otherwise) and quenched with 150 uL gDNA cell lysis buffer, 3 uL RNase A and 1 uL ProtK all from the Monarch gDNA extraction kit (NEB, T3010). gDNA was extracted following the extraction kit’s protocol with the exception that 600 uL of gDNA binding buffer were added to 400 uL quenched reaction and added to the extraction column on two spins of 2 minutes at 1000g. DNA was quantified using Qubit 1x dsDNA HS Assay Kit on a Qubit Flex Fluorometer.

### Enzymatic conversion

Methylated gDNA was digested using XbaI and AccI (NEB) to liberate a fragment containing the entire intact reporter sequence, and a 2-sided SPRI (0.4X followed by 1.8X) was performed to enrich for these fragments. DNA was converted using the Enzymatic Methyl-seq conversion module (NEB) following manufacturer’s instructions with slight modifications. First, 500 ng DNA (normally 200 ng) in 18 ul (normally 28 ul) was used as input to the oxidation reaction, where it was combined with 10 ul reconstituted TET2 reaction buffer + supplement, 1 ul oxidation supplement, 1 ul DTT, 3 ul oxidation enhancer (normally 1 ul) and 12 ul TET2 (normally 4 ul). This was mixed with 5 ul 0.4 mM Fe(II) solution and incubated for 1 hr at 37 °C, after which 1 ul Stop reagent (ProtK) was added and incubated for 30 min at 37 °C. 50 ul oxidized DNA was cleaned with 90 ul AmpureXP beads (1.8X), denatured by boiling in formamide at 90 °C (normally 85 °C) for 10 minutes, and quenched on ice. Deamination was performed by adding 10 ul APOBEC reaction buffer, 1 ul BSA, and 1 ul APOBEC, bringing the volume to 100 ul with H2O, and incubating at 37 °C for 3 hr. 100 ul DNA was cleaned with 100 ul AmpureXP beads (1X) before library preparation.

### Amplicon library preparation and sequencing

Amplicon library prep was performed in three rounds of PCR: 1) amplification out of the converted DNA 2) addition of part of R1 and R2 sequencing primers and 3) addition of remaining sequencing handles and sample indexes. For each 50 uL PCR1 reaction, we used 20 uL converted DNA, 2.5 uL each 10 uM forward and reverse primers (cTF223, cTF218 - see Supp. Table 2) and 25 uL NEBNext Q5U Master Mix (NEB, M0597) and ran the following thermocycling protocol: 30 seconds at 98 °C, then 21 cycles of 98 °C for 10 s, 61 °C for 30 s, 65 °C for 60 s, and then a final step of 65 °C for 5 minutes. PCR 1 product was then cleaned with a 0.9x AmpureXP bead clean and eluted in 21 uL H2O. For each 20 uL PCR2 reaction, we used 1 uL cleaned PCR 1 product, 2.5 uL each 10 uM forward and reverse primers (cTF401/402/, cTF403/404 - see Supp. Table 2) and 10 uL NEBNext Ultra Q5 Master Mix and ran the following thermocycling protocol: 30 seconds at 98 °C, then 6 cycles of 98 °C for 10 s, 66 °C for 30 s, 72 °C for 45 s, and then a final step of 72 °C for 5 minutes. Note that a mix of forward and reverse primers were used for PCR2, one set introduces an additional A after R1/R2 to improve sequence complexity. PCR 2 product was then cleaned with a 0.9x AmpureXP bead clean and eluted in 21 uL H2O. For each 25 uL PCR3 reaction, we used 20 uL cleaned PCR 2 product, 5 uL pooled 5 uM each forward and reverse indexing primers (oID1-7, oID8-24 - see Supp. Table 2) and 25 uL NEBNext Ultra Q5 Master Mix and ran the following thermocycling protocol: 30 seconds at 98 °C, then 6 cycles of 98 °C for 10 s, 68 °C for 30 s, 72 °C for 45 s, and then a final step of 72 °C for 5 minutes. PCR 3 product was then cleaned with a 0.9x AmpureXP bead clean and eluted in 17 uL H2O. Libraries were then quantified using Qubit 1x dsDNA HS Assay Kit for concentration and D1000 ScreenTape (Agilent Technologies, 5067-5582) for length. Libraries were pooled with 15% PhiX Sequencing Control v3 (Illumina, FC-110-3001) and 15% genomic EM-seq libraries for complexity and sequenced with 374 cycles on R1, 235 cycles on R2, 8 cycles on I1, and 4 cycles on I2 and with a MiSeq Reagent Kit v3 (600-cycle) (Ilumina, MS-102-3003) on a MiSeq.

### Genome-wide library preparation and sequencing

For shallow, genome-wide methylation analysis, methylated gDNA was sonicated to an average length of 400 bp using a Covaris E220 Focused-Ultrasonicator and combined with the methylated pUC19 and the unmethylated lambda-phage DNA (NEB). End prep and adaptor ligation was performed using the Enzymatic Methyl-seq kit (NEB). Conversion was performed identically to the amplicons. Libraries were constructed by PCR with the index primers for the EM-seq kit (NEB) with NEBNext Q5U Master Mix (NEB) using the following thermocycling protocol: 30 seconds at 98 °C, then 6 cycles of 98 °C for 10 s, 62 °C for 30 s, 65 °C for 60 s, and then a final step of 72 °C for 5 minutes. Libraries were sequenced either by spiking into the amplicon libraries for complexity, or on their own on a Next-seq using standard primers with 2x36 bp reads.

### Genome-wide SMF analysis

Reads were first trimmed to 36 bp using fastx_trimmer (http://hannonlab.cshl.edu/fastx_toolkit/index.html) and then aligned to the hg38, lambda, and pUC19 genomes using bwameth^59^. Duplicates were identified and removed using Picard MarkDuplicates (https://broadinstitute.github.io/picard/). Methylation fraction across the genome was called using MethylDackel (https://github.com/dpryan79/MethylDackel) and was aggregated either by sequence context or genomic position using custom scripts (available at https://github.com/GreenleafLab/amplicon-smf).

### Synthetic construct SMF analysis

Reads were aligned to a custom index (built from the individual PCR amplicons) using bwameth^59^. The resulting BAM files were filtered for alignment quality and uniqueness, unconverted reads were removed (by looking at the conversion of all non-GpC Cs), and bulk methylation was computed using MethylDackel (https://github.com/dpryan79/MethylDackel). Custom scripts were used to generate the single-molecule matrices and perform other quality control analyses. A Snakemake pipeline with associated conda environments is available at https://github.com/GreenleafLab/amplicon-smf.

### Single-molecule state calling model

Single-molecule states were assigned to each read using a maximum-likelihood approach. First, all possible single-molecule states (for a given amplicon) were enumerated (and precomputed to speed up future computations). A single-molecule state represents the occupancy status of each TetO, as well as the position (start and end coordinates, snapped to the nearest GpC) of all nucleosomes covering the molecule. The full list of states was enumerated by taking the Castesian product of the power set of the TetOs (each can be bound or unbound) and all possible positions of as many nucleosomes will fit along the molecule (mandating that the nucleosomes start and end at GpCs, have at least one accessible GpC in a linker between any pair of two nucleosomes, and are between 110-140 bp in length, the full width at half maximum of the longest stretch of uninterrupted protection across all molecules). For each hypothetical state (TetO occupancy and nucleosome positioning tuple), the expected methylation signal was computed by determining whether each GpC is accessible, protected by a nucleosome, or protected by a TF. These GpC occupancy states were then converted into probabilities using different methylation probabilities (unobserved) for each of the three possible occupancy states. This results in a matrix of size (number of states) x (number of GpCs) representing the probability that GpC j is methylated in state i. This matrix is then converted into a matrix P representing the probability of observing a converted base after the EM-seq reactions using the methylated (pUC19) and unmethylated (lambda) conversion rates from the controls. The maximum likelihood state for each input molecule (from single-molecule methylation matrix M, which is (number of GpCs) x (number of molecules)) was computed using a Bernoulli likelihood across all GpC positions in vectorized format (i.e. argmax(log(P) * M + log(1-P) * (1-M))), resulting in a vector of the most likely state for each input molecule. The code is available at https://github.com/GreenleafLab/amplicon-smf/blob/master/workflow/scripts/classify_single_molecule_binding_v2.py.

### Partition function model

The equilibrium thermodynamic model was constructed by deriving a function that assigns each molecule an energy. This energy function takes as input every possible single-molecule configuration from the state-calling model and counts the number of TFs and nucleosomes occupying the enhancer. Nucleosome dyads must overlap the enhancer to be counted. Each bound TF or nucleosome contributes a binding energy to the total energy of the molecule, except that nucleosomes bound to molecules with TFs also bound have their affinity reduced. Thus, three parameters (E_TF_, E_nuc_, E_remodel_) are used to assign each molecule an energy. A given set of these parameters determines an energy, and therefore a Boltzmann probability, for each molecular state. We use maximum likelihood estimation (using scipy.optimize.minimize() with method=’Nelder-Mead’^60^) to fit the three parameters that optimize the probability of seeing all observed molecules.

Alternative models encoding TF-only cooperativity (i.e. not dependent on nucleosomes) were implemented by adjusting the energy function per molecule. In particular, we tested 3 alternate models: a thresholding model (wherein TF binding affinity was increased if >1 TF was bound to the molecule), a TF-TF interaction model (wherein each pair of TFs bound to the molecule have an energy of interaction), and a cofactor scaffold model (wherein each bound TF has an energy of interaction with a cofactor that can be present or absent). Code is available at https://github.com/GreenleafLab/amplicon-smf/blob/master/workflow/scripts/fit_partition_function_model_v3.py.

To model the effective rTetR-VP48 concentration as a function of the dox concentration, we modified the standard binding isotherm. Specifically, we included a “leak” term (with units of concentration) to account for the fact that this version of the rTetR DNA-binding domain still binds DNA in the absence of dox: 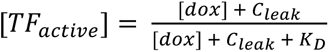. To compute the active TF fraction, we first note that the apparent TF binding energy from the thermodynamic model (*ΔE_TF_*,*_conc_*) depends on both the actual affinity and the TF concentration: 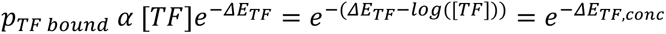. We fit the partition function model separately on each dox concentration and compute the fractional activity as a percent of the value at maximum dox concentration: 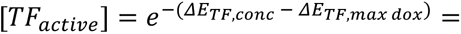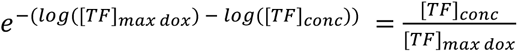. We then fit the apparent *C_leak_* and *K_D_* using scipy.optimize.curve_fit().

### Additive activation model

The equilibrium kinetic model of promoter activation, in which TFs act independently to activate the promoter (see **Fig. 3F**), was derived as follows with the assumption that the system is at steady state by 24 hours (where Prom_on_ = fraction of promoters active, Prom_off_ = fraction of promoters inactive = 1 - Prom_on_, k_on_ = promoter on rate, k_off_ = promoter off rate):

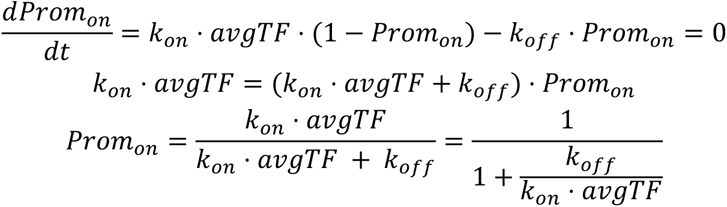

One parameter, representing k_off_/k_on_, was then fit to the relationship between average TF occupancy and the fraction of promoters active using scipy.optimize.curve_fit() and standard deviation on the parameter was calculated from the output covariance matrix, pcov, as follows: numpy.sqrt(numpy.diag(pcov)). Alternative molecular models (**see Fig. S5A**) were implemented with the same method and modifications to the on rate (cooperative: *k_on_* · *avgTF^n^*, thresholding: *k_on_* · *frac*_>=1*TF*_).

### Kinetic model

The kinetic model of TF binding, promoter activation, transcription, and translation (**see Fig. 5D**) was implemented as a continuous function for TF binding and as an ODE for all following steps. TF binding was modeled as a simple kinetic process that follows an exponential curve and is described by a single half-life (t_1/2_, see Equation 1). Note that t_1/2_ is related to TF:DNA on and off rates by the following relationship, 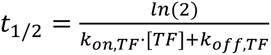. For rTetR-VP48, since there is a significant amount of binding that happens on ice during sample processing (see Fig. S5A), the sample of 0 hours induction with dox present in buffers during handling was used as the zero-time sample. The model was fit on background 0 for rTetR-VP48. The equation was then fit to average TF occupancy data over time, normalized to its maximum values (48 hours for rTetR-VP48, 6 hours for ISGF3), for amplicons with measurable binding (5-8 TetO, and 3-6 ISRE). The fit was performed using scipy.optimize.curvefit() and standard deviation on parameters was calculated from the output covariance matrix (numpy.sqrt(numpy.diag(pcov))). Promoter activation (dProm_on_/dt) was modeled using the non-steady-state version of the Additive Activation model (see above and Equation 2) and a two-parameter fit of k_on_ and k_off_. Transcription of RNA from active promoter states (dRNA/dt) was described by two parameters k_trs_ and k_decay,rna_, representing the combined effect of RNA loss through degradation and dilution (Equation 3). Similarly, translation of protein from RNA (dProt/dt) was described by two parameters k_trl_ and k_decay,prot_ (Equation 4). The combined ODE describing promoter activation, transcription and translation was fit across all amplicons using scipy.optimize.curvefit() which in turn called scipy.integrate.odeint() and passed in the TF_off,max_ for each amplicon; standard deviation on parameters was calculated from the output covariance matrix (numpy.sqrt(numpy.diag(pcov))).

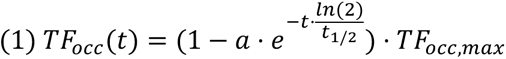

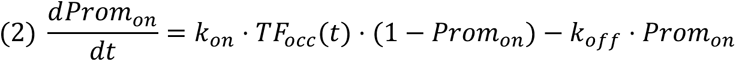

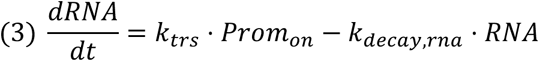

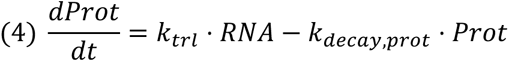

### Protein purification

TetON-3G (rTetR) was affinity purified with a C-terminal 6xHis-tag, where rTetR-HALO-TEV-6xHIS was expressed from a modified pD861 (ATUM) plasmid vector (pJS021) in T7-expression competent *E. coli* (NEB, C30131). 10 mL overnight cultures were diluted into 1 mL LB medium with 50 µg/mL kanamycin (Thermo Fisher, J60668.06) and grown at 37 °C and 120 RPM agitation. The expression of rTetR was induced at OD_600_=0.5-0.8 by adding 0.2% weight/volume L-rhamnose (Thermo Fisher, A16166.14) and expressed for 4 hours. Cultures were spun down at room temperature at 4600g for 40 minutes followed by 5000g for 40 minutes. Pellets were resuspended in 10 mL of buffer A (50 mM NaH2PO4, 300 mM NaCl, 20 mM imidazole, pH 8.0, one protease inhibitor tablet (Pierce, A32953)) and lysed on ice for 40 minutes with 10 µg/mL lysozyme (Thermo Fisher, 90082) and 25 units/mL benzonase I (Sigma Aldrich, E1014-25KU). Lysate was spun down at 14,000g for 30 min and supernatant was applied to Ni-NTA fast kit columns (Qiagen, 30600). Protein was washed with 16 mL buffer B (20 mM Hepes pH 7.3, 400 mM NaCl, 20 mM imidazole, 0.5 mM DTT) and eluted with buffer C (20 mM Hepes pH 7.3, 400 mM NaCl, 250 mM imidazole, 0.5 mM DTT). Eluted protein was then buffer exchanged into buffer D (20 mM Hepes pH 7.3, 400 mM NaCl, 0.5 mM DTT) on 10 kDa cut-off Amicon Ultra-0.5 centrifugal filters (Millipore Sigma, UFC901008). Protein was then further purified with a HiTrap Heparin HP column (Cytiva, 17-5247-01). The column was first washed with 5 mL 20% ethanol and then 5 mL buffer E (20 mM Hepes pH 7.3, 2 M NaCl, 0.5 mM DTT) before equilibrating with 2.5 mL buffer F (20 mM Hepes pH 7.3, 0.5 mM DTT). Protein was applied to the column, washed with 10 mL buffer F and eluted with increasing concentration of salt up to 2M NaCl in Buffer F (rTetR eluted around 200 mM NaCl and was >90% purified). Protein was then buffer exchanged into buffer D on 10 kDa cut-off Amicon Ultra-0.5 centrifugal filters and diluted to 50% glycerol. Protein was then flash frozen in liquid nitrogen and stored at - 80 °C for later use.

### EMSA

60 bp dsDNA with and without a TetO site was ordered as single-stranded and PAGE-purified oligos from IDT and formed from complementary strands by mixing 15 uL of 100 uM of each strand, 10 uL 10X PBS, 1 uL 1 M MgCl2, and 39 uL H2O and annealing at 94 °C for 3 minutes followed by 57 cycles of 50 seconds each where each cycle decreases the temperature 1 °C. Protein was incubated with DNA for one hour at room temperature in EMSA buffer (60% glycerol, 150 mM Hepes pH 7.3, 150 mM NaCl, 12 mM EDTA, 60 mM MgCl2, 0.5 mM DTT). Samples were loaded onto Mini-PROTEAN precast TBE gels (Bio-Rad, 4565013) in 1x TBE (Bio-Rad, 1610733) with TriDye Ultra Low Range DNA ladder (NEB, N0558S) and run at 100V for ∼50 minutes. Gels were then stained by shaking for 20 minutes in 50 mL TBE and 10,000x SYBR-green (Fisher Scientific, E33075) and washed with water. Gels were imaged with 4x bluegain and 120 second exposure, quantified using ImageJ, and fit with a binding isotherm 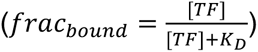 using scipy.optimize.curve_fit() and standard deviation on parameters was calculated from the covariance matrix (numpy.sqrt(numpy.diag(pcov))).

### ATAC-seq experiments and analysis

Bulk omni-ATAC was performed as in^61^ with 50,000 cells and two biological replicates per sample. Libraries were sequenced 2x36 with an NextSeq 500/550 High Output v2.5 75 cycles kit (Illumina, 20024906) on a NextSeq 550. Bulk ATAC-seq data processing of fastq files was performed with snakeATAC (https://github.com/GreenleafLab/snakeATAC_singularity) and alignment to hg38.

To compare SMF and ATAC data, ATAC peaks were first standardized to 200 bp wide and grouped by signal into 100 quantile bins. Average ATAC coverage was computed per bin. Methylation signal from SMF was computed by averaging over all GpCs in all reads overlapping the ATAC peaks and averaged by quantile.

Visualization of ATAC peaks at ISGs was performed by loading BigWig files with 100 bp windows into the Integrative Genomics Viewer^62^. BAM files were then further processed with ChrAccR (https://github.com/GreenleafLab/ChrAccR) to perform chromVar^63^ (IFN_ATAC_chraccr.rmd) and ATAC footprinting analyses. ATAC footprinting of ISREs was performed on genome-wide matches to Vierstra Non-redundant TF motif clustering (V2.1 BETA-HUMAN)^1^ cluster AC0188 IRF/STAT|IRF using ChrAccR function getMotifFootprints and RPM and Tn5 bias normalization (IFN_ATAC_footprinting.R). Computing average ATAC signal around IFN promoters, a list of hg38 TSS was extended by 250 bp in either direction, and average ATAC-seq coverage was computed over these peaks using bedtools coverage (vX). Signal was RPM normalized and averaged across all ISGs, which were defined as genes with >2-fold increase in expression 24 hours after IFN-β stimulation (as measured by bulk RNA-seq, see below) and which contained ≥3 ISRE motifs in the promoter. A null set of genes was selected by choosing the expressed gene with the closest TPM to each ISG pre-stimulation.

### Bulk RNA-seq of IFN-treated cells and analysis

K562 cells were cultured in 12-well plates and treated with 10 ng/mL interferon beta (PeproTech, #300-02BC) for 24 hours, after which 1 x 10^6^ cells were harvested by centrifugation at 300 x g for 5 minutes. Cells were lysed and homogenized using QIAshredder columns (Qiagen #79654) and RNA was extracted with the RNeasy Mini Kit (Qiagen #74104). The RNA Integrity Number (RIN) for all samples was 10 as assessed by an RNA Nano Kit (Agilent #5067-1511) on an Agilent Bioanalyzer, provided by the Stanford Protein and Nucleic ACID (PAN) Biotechnology Facility. 500 ng of purified RNA for each sample was enriched for polyadenylated mRNA with the NEBNext Poly(A) mRNA Magnetic Isolation Module (NEB #E7490) and then used as input for the NEBNext Ultra II RNA Library Prep Kit (NEB #E7770S). Sample preparation was performed as per manufacturer’s specifications, with nine PCR cycles used for final library amplifications. Library size distributions were confirmed using the High Sensitivity DNA Kit (Agilent #5067-4626) on an Agilent Bioanalyzer. Sample concentrations were measured using the Qubit dsDNA HS Assay Kit on a Qubit 4 Fluorometer, and all samples were pooled at equimolar ratios and sequenced on a NextSeq 550 with 2 x 37 cycles.

Sequencing reads were demultiplexed with bcl2fastq. Hisat2-build was used to build a reference transcriptome with a FASTA of the GRCh38 human reference genome and the accompanying GTF genome annotation file, and hisat2 was used to align the paired reads to the reference. Output SAM files were converted to BAM files using samtools and differential expression analysis was performed in R with the Bioconductor DESeq2 package^64^ using a set of custom R scripts (IFN_RNA_processing.R) that were largely based on the workflow and commands described in the following tutorial: http://bioconductor.org/help/course-materials/2016/CSAMA/lab-3-rnaseq/rnaseq_gene_CSAMA2016.pdf.

### ChIP-seq experiments

ChIP-seq was performed as previously described^65^ with modifications. For each ChIP reaction, 2x10^7 cells were used as input together with a spike-in of mouse chromatin used for orthogonal normalization. Cells were crosslinked with 1% formaldehyde for 15 min at room temperature, followed by quenching in glycine (final concentration 0.125 M). Cells were then centrifuged, resuspended in 1xPBS, centrifuged again, and stored at -80C. On the first day of the ChIP procedure, 10 uL Protein A Dynabeads (Thermofisher 10002D) were added to DNA Lo-Bind tubes and washed three times on a magnetic rack with 1 mL mg/mL BSA. Beads were resuspended in 1 mL BSA, 5 ug anti-H3K27ac antibody (Abcam ab4729) were added, and the antibodies were coupled to the beads overnight on a rotor at 4 °C. On the second day, crosslinked chromatin was resuspended in 1 mL Farnham Lysis Buffer (FLB; 5 mM HEPES pH 8.0, 85 mM KCl, 0.5% NP-40/IGEPAL, Roche Protease Inhibitor Cocktail), centrifuged, then resuspended in 1 mL FLB and centrifuged again, then resuspended in 880 uL RIPA buffer (1x PBS, 1% NP-40, 0.5% Sodium deoxycholate, 0.1% SGS, Roche protease inhibitor cocktail), and sheared using Covaris E220 Focused-Ultrasonicator. Beads were washed again three times with 1 mL BSA on a magnetic rack, resuspended in 100 uL BSA, and the chromatin in RIPA buffer was added to them, then incubated overnight at 4 °C on a rotor. On the third day, beads were washed five times with LiCl buffer (10mM Tris pH 7.5, 500 mM LiCl, 1% NP-40, 1% Sodium deoxycholate) for 10 minutes on a rotator at 4 °C followed by removal of the buffer using a magnetic rack, then washed once with 1 mL 1x TE buffer, and resuspended in 200 uL IP elution buffer (1% SDS, 0.1 M NaHCO_3_). Chromatin was eluted off the beads by incubation at 65 °C followed by centrifugation at max speed for 3 minutes, and transfer to fresh DNA Lo-Bind tubes. Crosslinks were reversed by addition of 2 uL Proteinase K (Promega) and incubation at 65 °C for 12-16 hours in a Thermomixer. DNA was purified by adding an equal volume (200 uL) 25:24:1 Phenol:Chloroform:Isoamyl alcohol, vortexing, and centrifugation for 3 minutes at max speed. The top phase was then purified using the MinElute kit (Qiagen), eluting in 50 uL 55°C EB buffer. Sequencing libraries were prepared using the NEBNext Ultra II kit (NEB E7645) following the manufacturer’s instructions. Sequencing was performed on a NextSeq in 2x38 bp format.

### ChIP-seq data processing

Sequencing reads were aligned against a custom genome index containing the hg38 version of the human genome, the reporter sequence, and the mm10 version of the mouse genome using Bowtie (version 1.0.1)^66^ with the following settings: “-v 2 -k 2 -m 1 -t --best --strata -q X 1000”. Duplicate alignments were removed using the MarkDuplicates function in PicardTools (version 1.99). Peaks were called using MACS2 (version 2.1.0)^67^ with default settings (“-g hs”). Additional analysis was carried out using custom Python scripts (https://github.com/georgimarinov/GeorgiScripts)^68^. Coverage over the reporter was normalized using the total number of reads mapped to the spike-in mouse DNA prior to RPM normalization.

## Data availability

All high-throughput sequencing datasets generated in this study are available in the NCBI Sequencing Read Archive (BioProject PRJNA1071686). Any additional information required to reanalyze the reported data in this paper is available from the lead contact upon request.

## Code availability

The synthetic SMF analysis and state-calling software are available on GitHub (https://github.com/GreenleafLab/amplicon-smf). The ATAC-seq analysis is available on GitHub (https://github.com/GreenleafLab/snakeATAC_singularity) and additional .ipynb and R scripts for processing RNA-seq and ATAC-seq are available in Supplementary Information. Any further custom code for computational analyses and visualization are available from authors upon request.

## Supporting information

Supplemental Plasmid Maps

## Acknowledgements

We thank Betty Liu for Tn5 and help with ATAC-seq and processing, Soon il Higashino and Sage Allen for keeping our labs running, Surag Nair for designing background sequence #2, Connor Ludwig for pCL056 and help with RNA-seq, Eyal Metzl-Raz for the coffee corner, and all the members of the Greenleaf, Bintu, and Engreitz laboratories for helpful discussions and feedback. This work was supported by grant numbers NSF GRFP DGE-1656518 (B.R.D., M.M.H., and J.M.S.) and DGE-2146755 (C.R.M. and Y.T.), Stanford Interdisciplinary Graduate Fellowship affiliated with Stanford Bio-X (B.R.D., J.M.S.), Stanford Bio-X Bowes Fellowship (A.R.T.), Sarafan Chem-H Chemistry-Biology Interface Training Grant (A.R.T.), NIH Training Program grant T32GM145402 (A.R.T.), Stanford Graduate Fellowship (M.M.H.), and Stanford VPGE EDGE Fellowship (M.M.H. and C.R.M.). E.M. acknowledges support from the Swedish Research Council (grant 2020-06459), the Foundation Blanceflor, and the Science for Life Laboratory (SciLifeLab). This work was supported by NIH grants UM1HG009436 and P50HG007735 (to W.J.G.), and NIH-NIGMS R35M128947 (to L.B.). W.J.G. was a Chan Zuckerberg Biohub investigator and acknowledges grants 2017-174468 and 2018-182817 from the Chan Zuckerberg Initiative.

## Supplementary Information

Tables:

1. TableS1: Oligo libraries and DNA sequences installed at the reporter.
2. TableS2: Primers used for targeted amplification for sequencing and for library cloning into the reporter.
3. TableS3: Summary metrics (e.g. average occupancy, fraction of promoters active) for all SMF experiments.
4. TableS4: Single-molecule state model calls for all SMF experiments.
5. TableS5: Flow cytometry summary metrics for all experiments.
6. TableS6: Citrine RNA RT-qPCR data for all experiments.

Plasmid maps:

1. pCL056.gb: AAVS1 PuroR-RECloneSite-minCMV-IGKleader-hIgG1_FC-Myc-PDGFRb-T2A-Citrine-PolyA
2. pMMH006.2.gb: piggyBac TRE3G-hOct4_DBD-T2A-mCherry-pEF-rTetR(3G)-VP48-T2A-HygR
3. pMMH107.gb: piggyBac TRE3G-hSox2_DBD-T2A-mCherry-pEF-rTetR(3G)-T2A-HygR
4. pMMH090.gb: piggyBac pEF-rTetR(G72P)-3xFLAG-VP48-pEFcore-LTR-mCh-BSD
5. pMMH096.gb: piggyBac pEF-rTetR(G72P)-3xFLAG-TypeIIS-pEFcore-LTR-mCh-BSD
6. pJS021.gb: pRha-rTetR-HALO-TEV-6xHIS-Ori_pUC-kan(Km^r^)

## Author Contributions

B.R.D., M.M.H., and J.M.S. contributed equally as co-first authors, are listed alphabetically by last name and have agreed that any author can be listed first in reporting this study. M.M.H., B.R.D., J.M.S., L.B., and W.J.G. conceived of the study. M.M.H., B.R.D., and J.M.S. designed, performed and analyzed all SMF experiments, with assistance from G.K.M. and C.R.M. M.M.H. and B.R.D. optimized the SMF assay. J.M.S. and M.M.H. designed and cloned all libraries, and M.M.H., J.M.S., A.R.T., and B.R.D. made cell lines and performed tissue culture. G.K.M. and M.M.H. performed and G.K.M. analyzed ChIP-seq. J.M.S., M.M.H., and A.R.T. performed and J.M.S. analyzed flow cytometry. B.R.D., J.M.S., and C.R.M. performed and analyzed ATAC-seq, with help from D.D. A.R.T. performed and analyzed the RNA-seq experiments. J.M.S. and E.M. purified rTetR and performed EMSAs. B.R.D. wrote the SMF processing code and probabilistic binding model, with significant input from G.K.M., B.E.P., and Y.T. B.R.D. and M.M.H. wrote the Simple Competition and Nucleosome Destabilization models, J.M.S. and M.M.H. wrote the Additive Activation model, and J.M.S. wrote the kinetic model, all with significant input from L.B. and W.J.G. J.M.S., B.R.D., L.B., and W.J.G. wrote the manuscript, with input from all authors. L.B. and W.J.G. jointly supervised the work.

## Supplementary Figures

**Figure S1.**
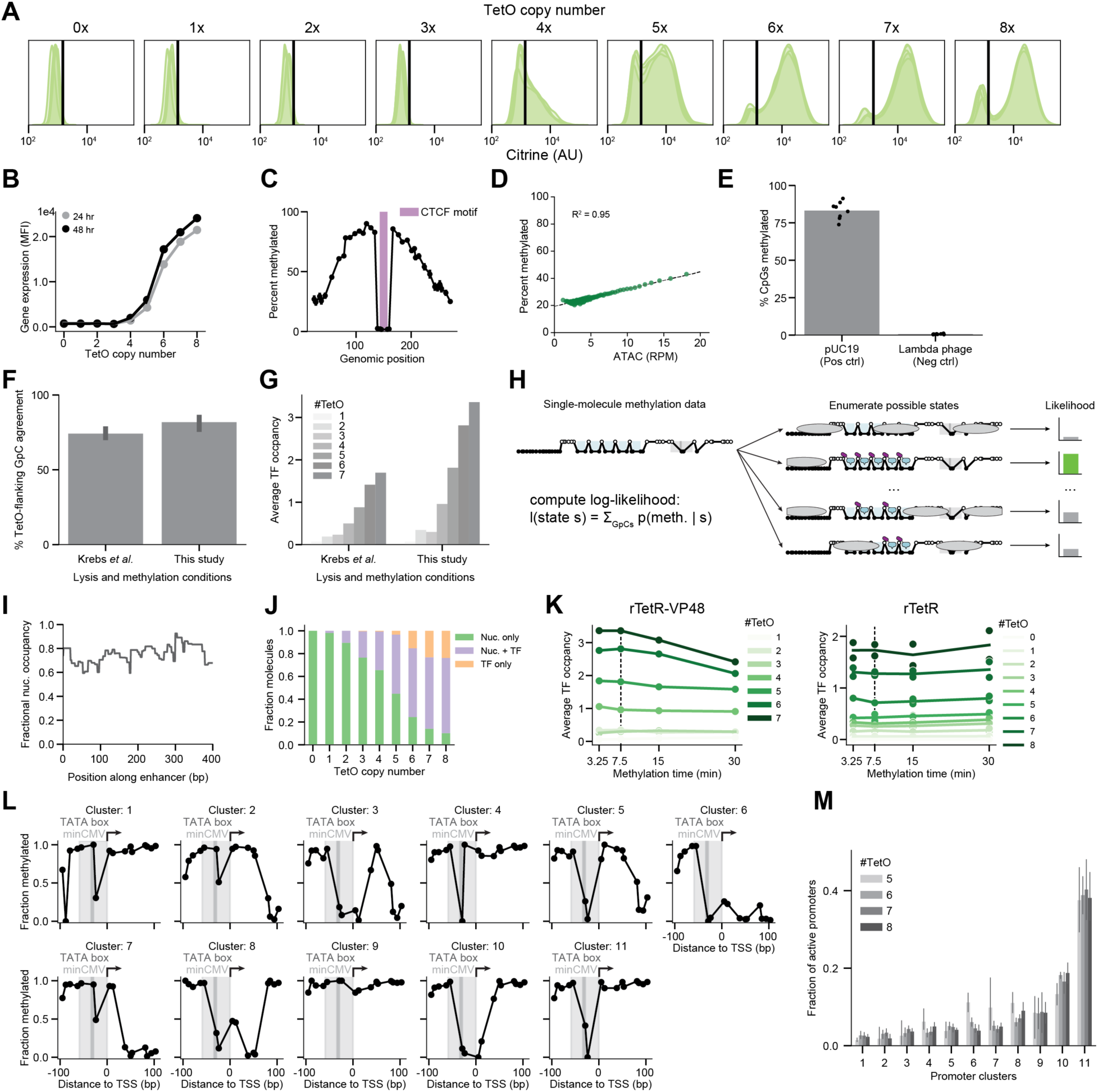
A) Flow cytometry distributions for Citrine fluorescence for the 0-8xTetO amplicons. Data are from 2-4 technical replicates. B) MFI for Citrine fluorescence for the 0x-8xTetO amplicons at 24 and 48 hours. C) Average methylation signal around a synthetic amplicon containing a single CTCF site (location marked in purple). Error bars represent the 95% CI from 3 biological replicates. D) Relationship between the average ATAC-seq signal (RPM) and average methylation probability from SMF for all ATAC-seq peaks genome-wide grouped into 100 quantile bins. E) Methylation probabilities at CpGs from the fully methylated (pUC19) and fully unmethylated (Lambda phage) EM-seq controls. Each dot represents an experiment and the bar represents the average. F) The percent of pairs of GpCs that flank the same TetO site that have the same methylation status on the same molecule. Error bars represent the 95% CI obtained by combining data across all pairs of GpCs that flank all TetO sites. G) Average TF occupancy across 0x-8xTetO amplicons under different lysis and methylation conditions. H) Cartoon schematic of the probabilistic binding model. For each observed single-molecule methylation signal, we enumerate all possible underlying molecular configurations of TFs and nucleosomes and assign each a likelihood of being observed. We then select the state with the maximum likelihood. I) The fraction of molecules with a nucleosome overlapping each position in the synthetic enhancer sequence. J) The fraction of molecules which have enhancers either a) completely covered by nucleosomes, b) only bound by TFs, or c) covered with a mix of the two as a function of the number of TetO sites. K) Average TF occupancy across all amplicons for both rTetR-VP48 and rTetR alone as a function of the duration of the methylation reaction. Dotted line represents the conditions used in our assay (7.5 min). L) Average methylation signal across the promoter region for molecules grouped into each of the 11 k-means clusters of “active” (nucleosome-free) promoters. M) Fraction of “active” (nucleosome-free) promoters that are found in each k-means cluster, faceted by the number of TetO sites in the amplicon. Error bars represent the 95% CI from 4 biological replicates. Cluster numbers are the same as in (L).

**Figure S2.**
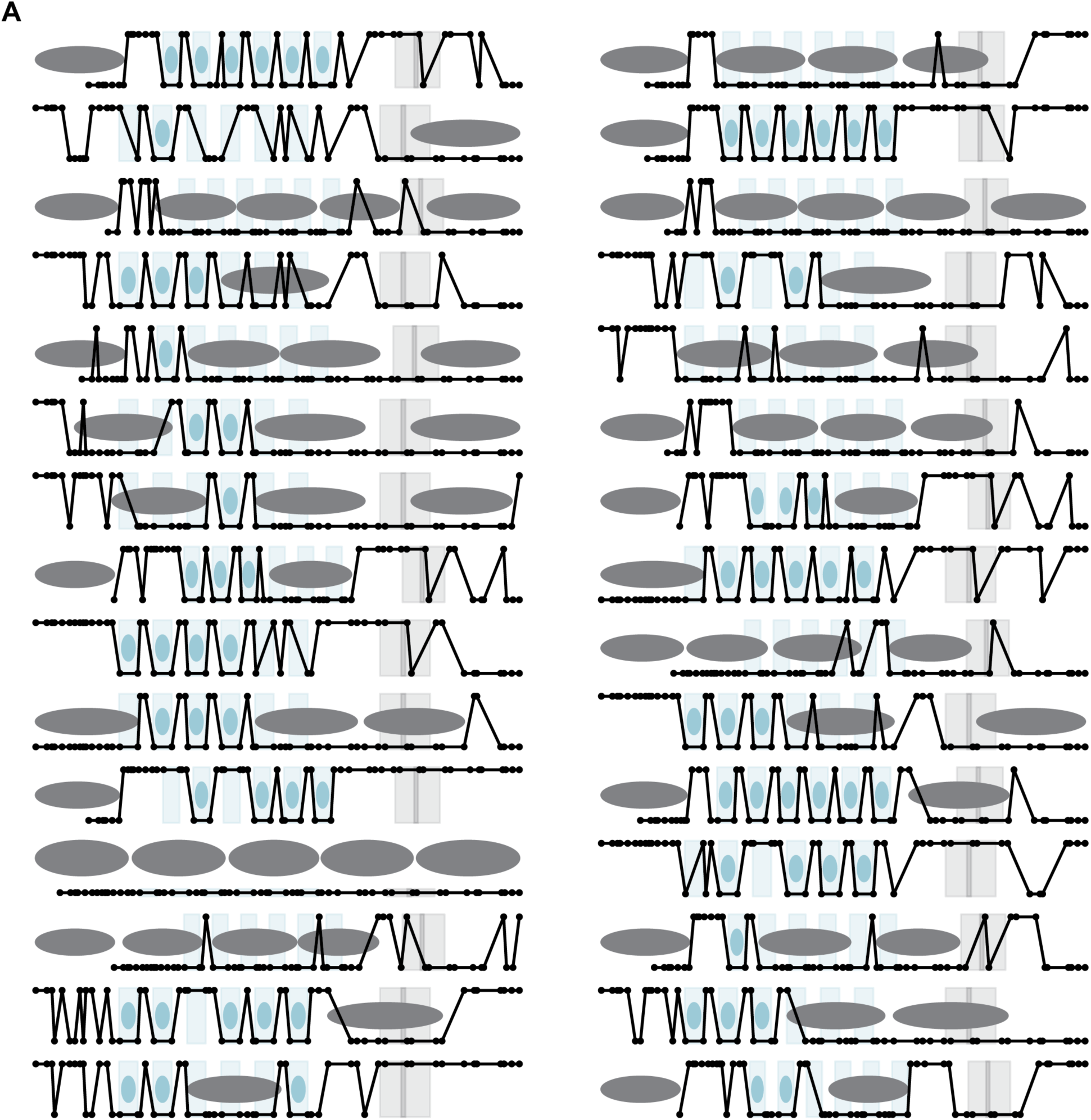
A) Exemplar data for 30 random single molecules, each of 6xTetO background 0, after 24 hours of dox induction (1,000 ng/ml) and binding model calls for nucleosomes (gray) and rTetR-VP48 (blue). For each molecule, points on the top are methylated (accessible) and points on the bottom are unmethylated (inaccessible).

**Figure S3.**
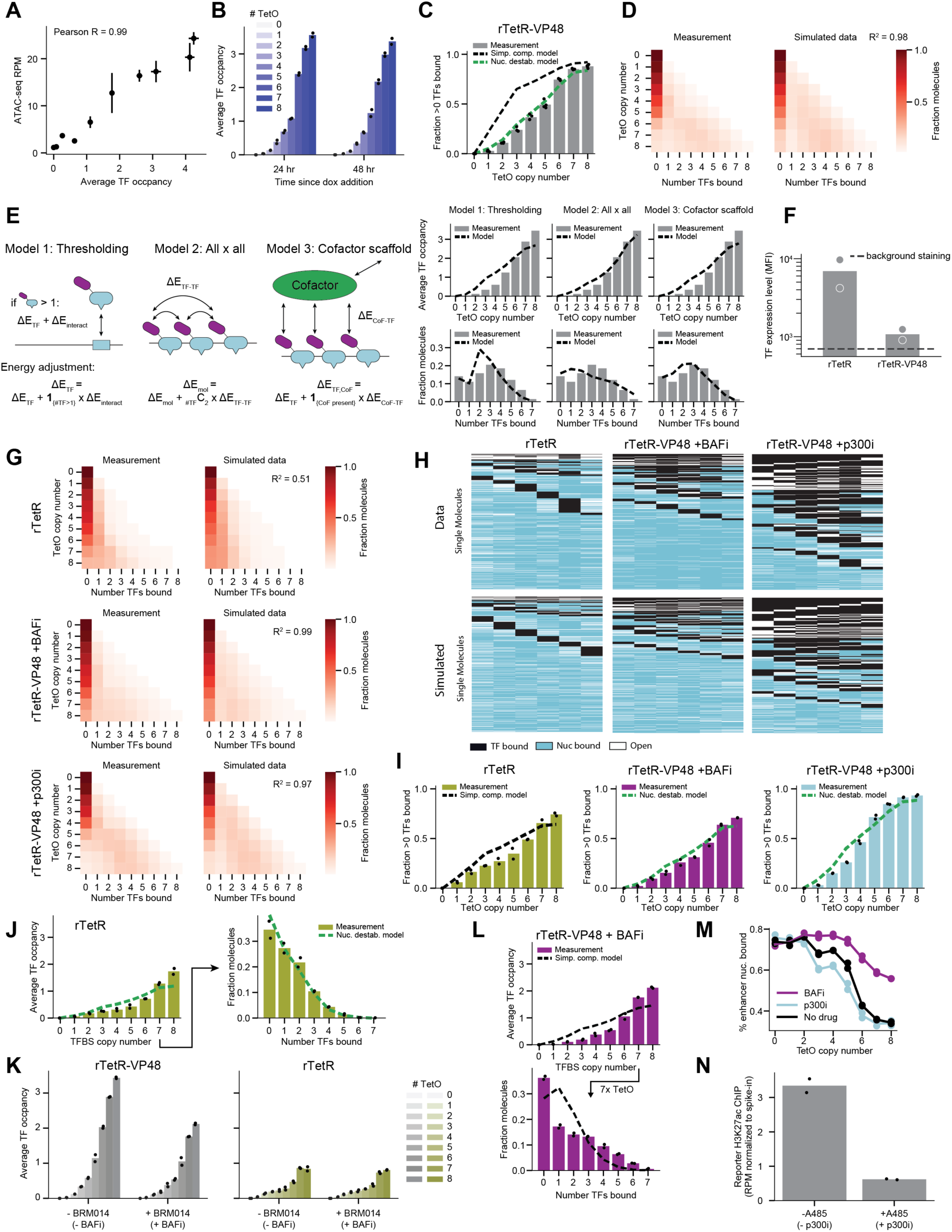
A) Relationship between average TF occupancy and ATAC-seq signal (RPM) for a selection of single enhancer amplicons from multiple backgrounds. Each dot represents a single enhancer. Error bars represent the 95% CI from 4 (SMF) or 2 (ATAC-seq) biological replicates. B) Average TF occupancy across 0x-8xTetO amplicons at 24 and 48 hours. C) Observed fraction of molecules with >0 TFs bound for enhancers with increasing number of TetO sites (four biological replicates) fit to either the Simple Competition (r^2^ = 0.44) or Nucleosome Destabilization (r^2^ = 0.99) models (from **Fig. 2D-E**). D) Observed occupancy distributions across all enhancers (left) with matching simulations from the Nucleosome Destabilization model (right). Each row represents a single enhancer and each column represents the number of TFs bound. The pixel intensity at (row i, column j) indicates the fraction of molecules with i TetO sites that have j TFs bound. Rows sum to 1. E) Three alternate TF cooperativity models (see Methods). Cartoons demonstrating how the TF cooperativity models are encoded (left) and fits to average occupancy and occupancy distribution data (right). F) Flow cytometry for FLAG-tagged TF expression for cells expressing rTetR or rTetR-VP48. Dotted line represents the level of background staining. Data are from two biological replicates. G) Observed occupancy distributions across all enhancers (left) with matching simulations from the Nucleosome Destabilization model (right) as in (D) for rTetR only, rTetR-VP48 treated with BAF inhibitor, and rTetR-VP48 treated with p300 inhibitor. H) Full molecular state representations for 10,000 measured (top) or simulated (bottom) molecules (as in Fig. 2G) for rTetR only, rTetR-VP48 treated with BAF inhibitor, and rTetR-VP48 treated with p300 inhibitor. Each column represents a TetO site and each row represents a molecule. Sites are colored by their occupancy status. I) Observed fraction of molecules with >0 TFs bound as in (C) for rTetR only, rTetR-VP48 treated with BAF inhibitor, and rTetR-VP48 treated with p300 inhibitor. Fits are either to the Simple Competition (rTetR only) or Nucleosome Destabilization (rTetR-VP48 + inhibitors) models (r^2^ = 0.86, 0.96, and 0.96, respectively). Data are from two biological replicates. J) Observed average occupancy (left) and occupancy distribution (right) of rTetR (two biological replicates) with a fit from the Nucleosome Displacement model (r^2^ = 0.83 and 0.96, respectively) (cf. **Fig. 2J-K**). K) Average TF occupancy across 0x-8xTetO amplicons for rTetR-VP48 (left) and rTetR only (right) with and without treatment with BAF inhibitor. Data are from two biological replicates. L) Observed average occupancy (top) and occupancy distribution (bottom) of rTetR-VP48 in the presence of the BAF inhibitor BRM014 (two biological replicates) with a fit from the Simple Competition model (r^2^ = 0.78 and 0.51, respectively) (cf. **Fig. 2L-M**). M) Observed fraction of molecules with nucleosomes overlapping the enhancer as a function of the number of TetO sites for rTetR-VP48, rTetR-VP48 treated with BAF inhibitor, and rTetR-VP48 treated with p300 inhibitor. N) Average H3K27ac signal across the synthetic reporter locus with and without p300 inhibition. Data are from two biological replicates, and the signal is normalized to spike-in mouse chromatin.

**Figure S4.**
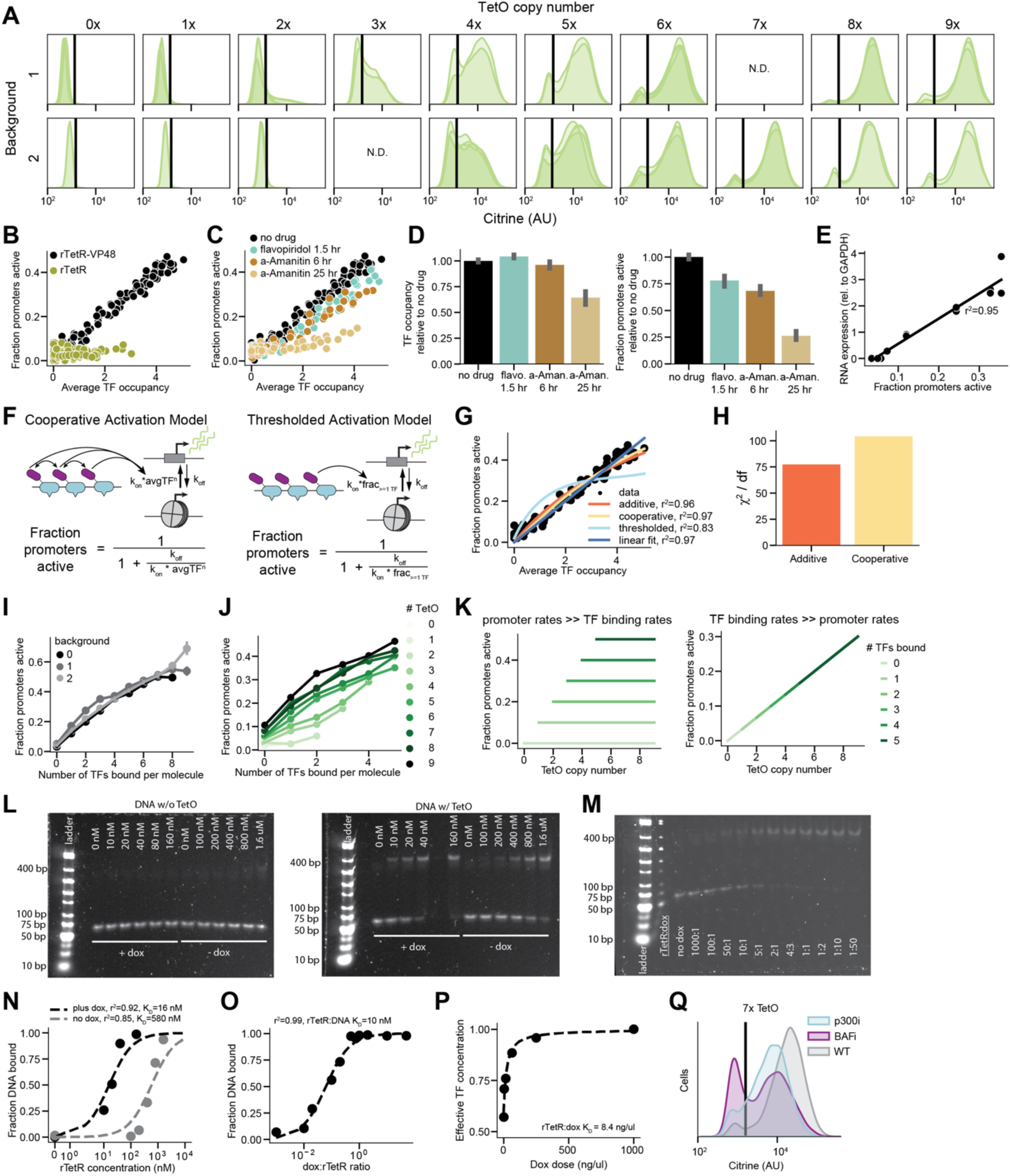
A) Flow cytometry distributions for the 0-9xTetO amplicons in backgrounds 1 and 2. Data are from 2-4 technical replicates. B) Relationship between rTetR-VP48 (black) and rTetR (green) occupancy and the fraction of active promoters. C) Relationship between rTetR-VP48 occupancy and the fraction of active promoters without drugs (black) and across transcription inhibition conditions with 24 hours of dox: flavopiridol for 1.5 hours (cyan) and alpha-amanitin for 6 hours (brown), alpha-amanitin for 25 hours (tan). D) Quantification of C for TF occupancy (left) and fraction of promoters active (right) across transcription inhibition conditions relative to no drug treatment for 5x-8xTetO. Alpha-amanitin for 25 hours (tan) likely reduces rTetR-VP48 concentration due to global reduction of transcription. E) Relationship between fraction of promoters active and RNA expression (measured by RT-qPCR, normalized to GAPDH levels) on 0-8xTetO with two technical replicates each. Black line is a linear fit. F) Schematics of alternative models relating TF binding to promoter activation. Cooperative Activation (left) assumes activation is cooperative with the number of TFs bound on average (kon*avgTF^n^). Thresholded Activation (right) assumes activation of the promoter occurs when there is at least 1 TF present (kon*frac>=1 TF). G) Model fits for additional models in F on the relationship between average rTetR-VP48 occupancy and fraction of promoters active. Fit parameters for cooperative model: kon/koff = 0.13 ± 0.006 TF^-1^, n = 1.1 ± 0.04. Fit parameter for thresholded model: kon/koff = 0.5 ± 0.1. H) Chi-squared per degree of freedom comparing Additive and Cooperative Activation model fits in C. I) Instantaneous relationship between the number of rTetR-VP48s bound and the promoter state on the same molecule for all TetO copy numbers across backgrounds. J) Instantaneous relationship between the number of rTetR-VP48s bound (up to 9) and the promoter state on the same molecule separating molecules by TetO copy number. K) Example plots to compare to data in Fig. 3H representing the expected relationship between TetO copy number, number of TFs bound, and promoter activity on the same molecules if promoter rates are much faster than TF binding rates (left) or vice versa (right). L) EMSA of rTetR (concentration noted above each lane) binding to 60 bp target DNA (1 nM) without a TetO site (left) and with a TetO site (right) in the presence and absence of doxycycline (1:50 rTetR to dox concentration). M) EMSA of rTetR (160 nM) binding to 60 bp target DNA (1 nM) with a TetO site across varying dox concentrations (ratio relative to rTetR concentration noted under each lane) (left). N) Quantification of EMSA gel with TetO sites in L in the presence (black) and absence (gray) of dox with binding isotherm fits (y=x/(KD+x)). O) Quantification of EMSA gel in M with binding isotherm fit (y=x/(KD+x)) where KD is the affinity of rTetR to DNA and the affinity of dox to rTetR is assumed to be much smaller. P) Quantification of the apparent TF concentration from relative *in vivo* rTetR-VP48 concentration (from binding energy in Nucleosome Destabilization model) across dox concentrations with binding isotherm fit. Q) Flow cytometry distributions for a 7xTetO reporter cell line without drugs (gray), under p300 inhibition (cyan) and under BAF inhibition (purple) after 24 hours of dox induction.

**Figure S5.**
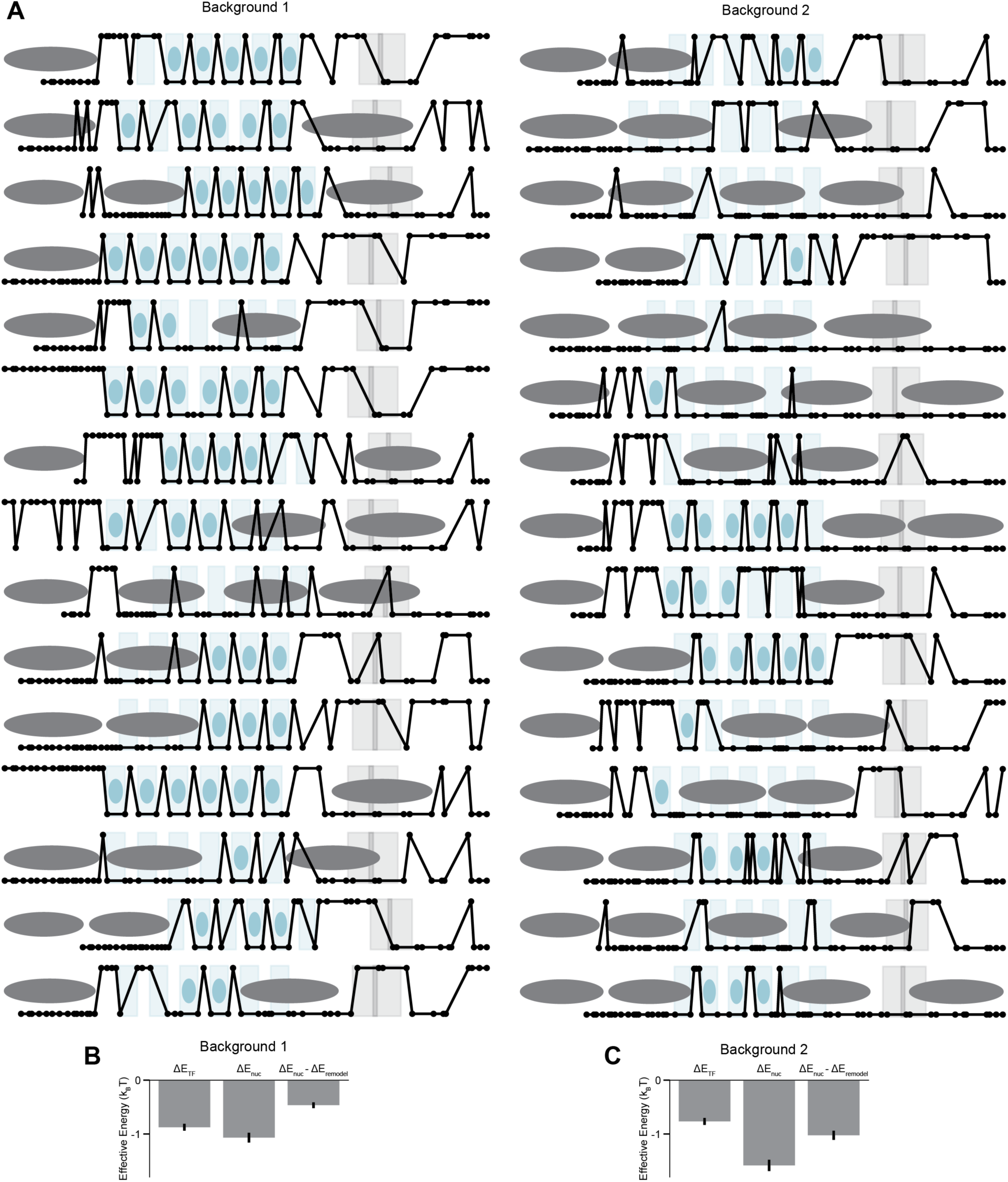
A) Exemplar data for 15 random single molecules, each of 6xTetO background 1 (left) and background 2 (right), 24 hours after dox induction (1000 ng/ml) and binding model calls for nucleosomes (gray) and rTetR-VP48 (blue). For each molecule, points on the top are methylated (accessible) and points on the bottom are unmethylated (protected). B-C) Model parameters for the equilibrium partition function model fit on background 1 and 2 molecules, respectively.

**Figure S6.**
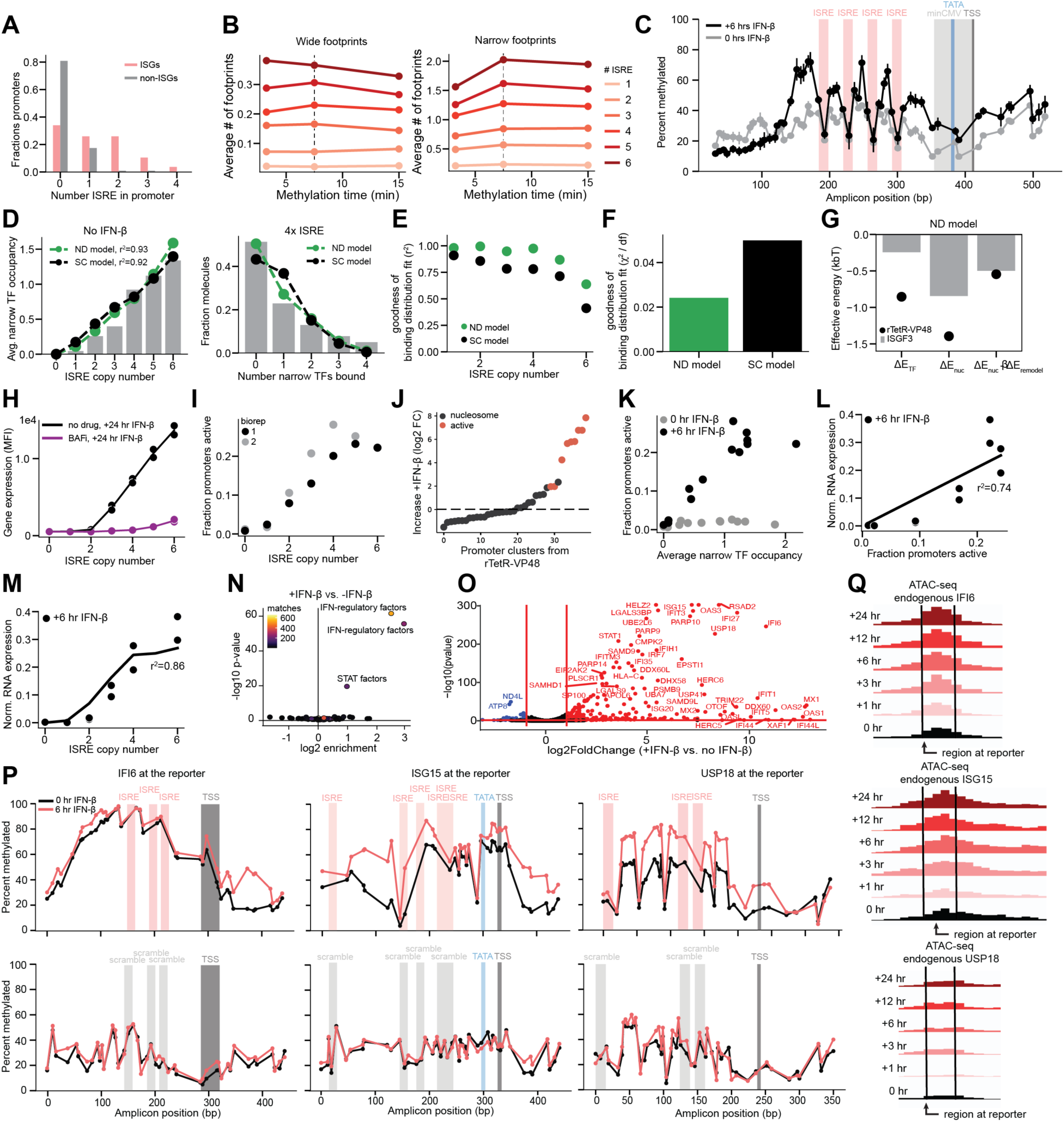
A) Distributions of the number of ISREs within a 500 base pair window around TSSs of ISGs identified from bulk RNA-seq (pink) and of non-ISGs (gray). B) Relationship between methylation time and measured average occupancy across amplicons after 24 hours of IFN-ꞵ stimulation for wide footprints (left) and narrow footprints (right). Dashed black line is the methylation time chosen for all subsequent experiments. C) Aggregated methylation data obtained for the construct with 4 ISRE sites present, with (black) and without (gray) IFN-ꞵ present. Error bars are standard deviation between two biological replicates. D) Relationship between ISRE copy number and average narrow footprint occupancy for the average of two biological replicates (left) and the distribution of narrow footprints bound for 4x ISRE sites (right) prior to stimulation fit by the Simple Competition model (black) and Nucleosome Displacement model (green). For both models, the nucleosome energy is fit to the nucleosome value measured in rTetR-VP48 experiments. E-F) Goodness of fit of number bound distributions in D by r^2^ across ISRE copy numbers (E) and chi-squared per degrees of freedom (F) on the number bound distributions for both models. F) Fit parameters (averaged across two biological replicates) from Nucleosome Displacement model (gray) in D; black dots are rTetR-VP48 parameters. G) Relationship between ISRE copy number and gene expression (as measured by flow cytometry at 24 hours of IFN-ꞵ stimulation) with (purple) and without (black) BAF inhibition for two technical replicates. H) Relationship between ISRE copy number and fraction of promoters active across two biological replicates at 6 hours of IFN-ꞵ stimulation. I) Enrichment of active promoter clusters (classified using rTetR-VP48 data) at 6 hours of IFN-ꞵ stimulation compared to 0 hours for two biological replicates. J) Relationship between narrow footprint occupancy and fraction of promoters active with (black) and without (gray) 6 hours of IFN-ꞵ stimulation. K) Relationship between fraction of promoters active and RNA expression (measured by normalized RT-qPCR) on 0-6xISRE for two technical replicates. Black line is a linear fit. L) Relationship between ISRE copy number and gene expression (as measured by RT-qPCR of Citrine mRNA) for two technical replicates. Black line is the coupled Additive Activation model and linear fit from fraction promoters active to gene expression with input of measured average wide TF occupancy. M) Differential ATAC-seq chromatin accessibility of TF motif types after 6 hours of IFN-ꞵ stimulation over two biological replicates. N) Differential RNA-seq expression after 6 hours of IFN-ꞵ stimulation over two biological replicates. O) Aggregated methylation data obtained for three reporter constructs with endogenous ISG promoters and proximal enhancers present (IFI6, ISG15, USP18) with intact ISREs (top) and scrambled ISREs (bottom) with (pink) and without (black) IFN-ꞵ present. P) ATAC-seq tracks of normalized chromatin accessibility to the same scale over an IFN-ꞵ stimulation timecourse for IFI6, ISG15, and USP18. Black lines denote the region that was installed at the reporter.

**Figure S7.**
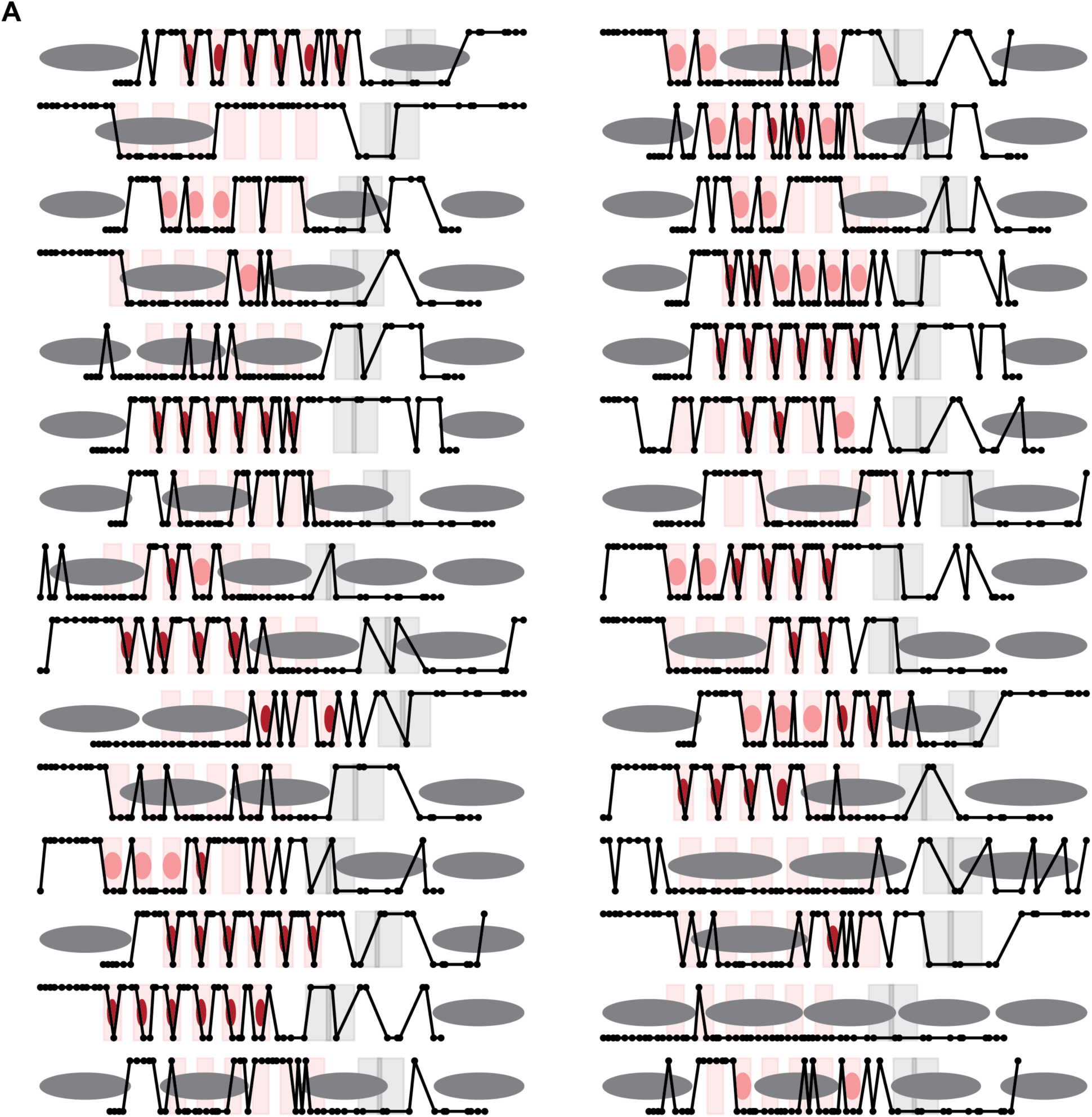
A) Exemplar data for 30 random single molecules, each with 6xISRE, after 6 hours of IFN-ꞵ and binding model calls for nucleosomes (gray), narrow footprints (red), and wide footprints (pink). For each molecule, points on the top are methylated (accessible) and points on the bottom are unmethylated (protected).

**Figure S8.**
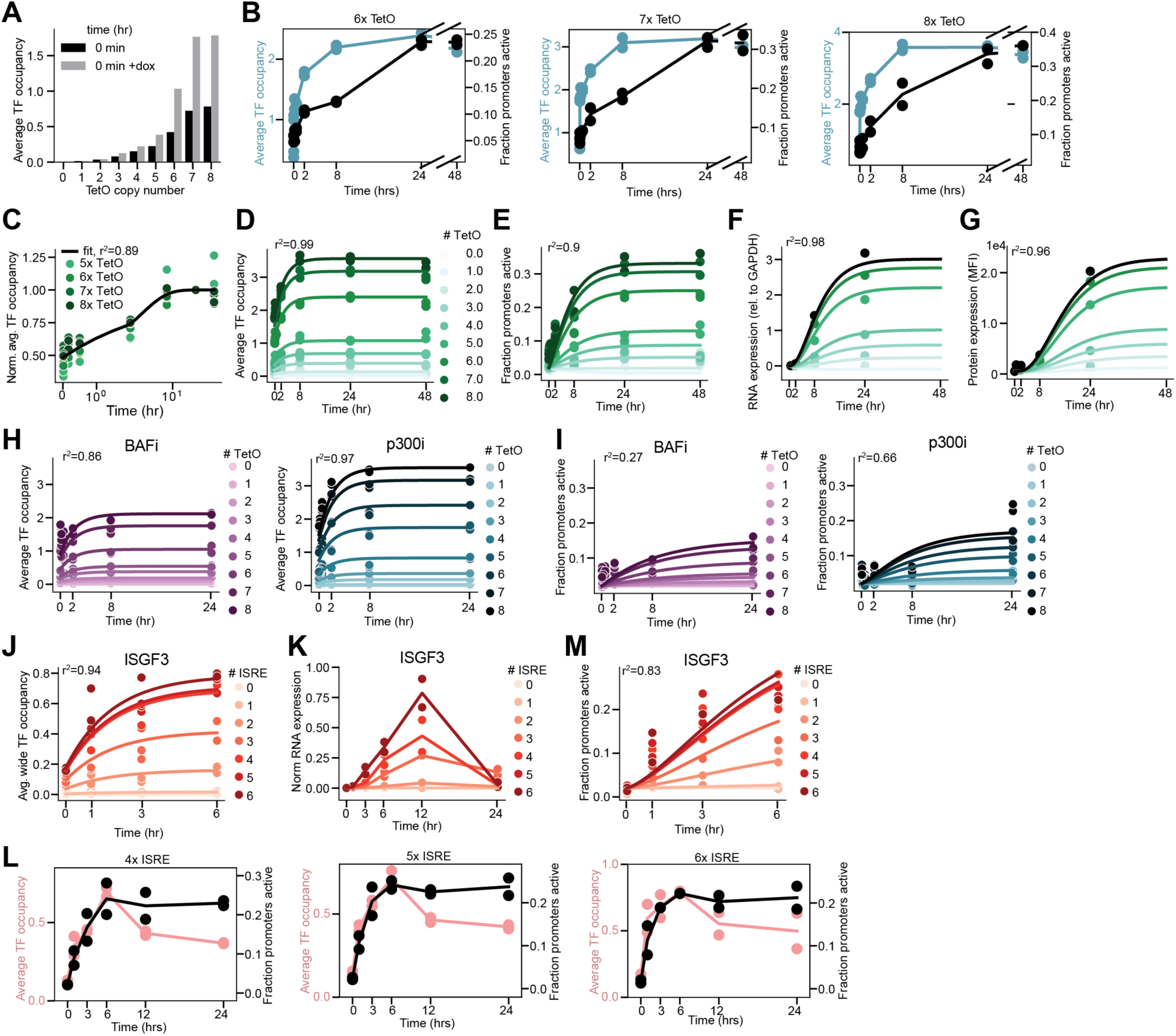
A) rTetR-VP48 occupancy without (black) and with dox (gray) present during sample processing across TetO copy numbers. B) Representative temporal delay between rTetR-VP48 occupancy (blue) and promoter activation (black) for two biological replicates of 6-8 TetO sites. C) Fit of single exponential to average rTetR-VP48 occupancy normalized to 48 hour value for two biological replicates of 5-8 TetO sites. D-G) Average rTetR-VP48 occupancy (D), fraction of promoters active (E), Citrine RNA expression as measured by RT-qPCR (F) and Citrine protein levels as measured by flow cytometry (G) over time for two biological replicates of 0-8 TetO sites. Lines are the fit kinetic model (as described in Fig. 5D). H-I) Average rTetR-VP48 occupancy (H) and fraction of promoters active (I) over time for two biological replicates of BAF inhibition (purple) and p300 inhibition (blue). Lines are the fit kinetic model (as described in Fig. 5D). K) Average wide footprint occupancy over time during the activation process (up to six hours) for two biological replicates of 0-6 ISRE sites. Lines are fit kinetic model (as described in Fig. 5D). L) Citrine RNA expression as measured by normalized RT-qPCR over time for two technical replicates of 0,1,2,3,4, and 6 ISRE sites. M) Representative lack of temporal delay between wide footprint occupancy (pink) and promoter activation (black) for two biological replicates of 4-6 ISRE sites. N) Fraction of promoters active over time during the activation process (up to six hours) for two biological replicates of 0-6 ISRE sites. Lines are fit kinetic model (as described in Fig. 5D).

## References

1. Vierstra, J. et al. Global reference mapping of human transcription factor footprints. Nature 583, 729–736 (2020).

2. Giniger, E. & Ptashne, M. Cooperative DNA binding of the yeast transcriptional activator GAL4. Proc. Natl. Acad. Sci. U. S. A. 85, 382–386 (1988).

3. Pettersson, M. & Schaffner, W. Synergistic activation of transcription by multiple binding sites for NF-kappa B even in absence of co-operative factor binding to DNA. J. Mol. Biol. 214, 373–380 (1990).

4. Spitz, F. & Furlong, E. E. M. Transcription factors: from enhancer binding to developmental control. Nat. Rev. Genet. 13, 613–626 (2012).

5. Thanos, D. & Maniatis, T. Virus induction of human IFN beta gene expression requires the assembly of an enhanceosome. Cell 83, 1091–1100 (1995).

6. Mirny, L. A. Nucleosome-mediated cooperativity between transcription factors. Proc. Natl. Acad. Sci. U. S. A. 107, 22534–22539 (2010).

7. Biddie, S. C. et al. Transcription factor AP1 potentiates chromatin accessibility and glucocorticoid receptor binding. Mol. Cell 43, 145–155 (2011).

8. Fryer, C. J. & Archer, T. K. Chromatin remodelling by the glucocorticoid receptor requires the BRG1 complex. Nature 393, 88–91 (1998).

9. Herschlag, D. & Johnson, F. B. Synergism in transcriptional activation: a kinetic view. Genes Dev. 7, 173–179 (1993).

10. Martinez-Corral, R. et al. Transcriptional kinetic synergy: A complex landscape revealed by integrating modeling and synthetic biology. Cell Syst 14, 324–339.e7 (2023).

11. Kelly, T. K. et al. Genome-wide mapping of nucleosome positioning and DNA methylation within individual DNA molecules. Genome Res. 22, 2497–2506 (2012).

12. Krebs, A. R. et al. Genome-wide Single-Molecule Footprinting Reveals High RNA Polymerase II Turnover at Paused Promoters. Mol. Cell 67, 411–422.e4 (2017).

13. Shipony, Z. et al. Long-range single-molecule mapping of chromatin accessibility in eukaryotes. Nat. Methods 17, 319–327 (2020).

14. Stergachis, A. B., Debo, B. M., Haugen, E., Churchman, L. S. & Stamatoyannopoulos, J. A. Single-molecule regulatory architectures captured by chromatin fiber sequencing. Science 368, 1449–1454 (2020).

15. Sönmezer, C. et al. Molecular Co-occupancy Identifies Transcription Factor Binding Cooperativity In Vivo. Mol. Cell 81, 255–267.e6 (2021).

16. Gossen, M. et al. Transcriptional activation by tetracyclines in mammalian cells. Science 268, 1766–1769 (1995).

17. Vaisvila, R. et al. Enzymatic methyl sequencing detects DNA methylation at single-base resolution from picograms of DNA. Genome Res. 31, 1280–1289 (2021).

18. Ackers, G. K., Johnson, A. D. & Shea, M. A. Quantitative model for gene regulation by lambda phage repressor. Proc. Natl. Acad. Sci. U. S. A. 79, 1129–1133 (1982).

19. Bintu, L. et al. Transcriptional regulation by the numbers: models. Curr. Opin. Genet. Dev. 15, 116–124 (2005).

20. Kim, H. D. & O’Shea, E. K. A quantitative model of transcription factor-activated gene expression. Nat. Struct. Mol. Biol. 15, 1192–1198 (2008).

21. Neely, K. E. et al. Activation domain-mediated targeting of the SWI/SNF complex to promoters stimulates transcription from nucleosome arrays. Mol. Cell 4, 649–655 (1999).

22. Yudkovsky, N., Logie, C., Hahn, S. & Peterson, C. L. Recruitment of the SWI/SNF chromatin remodeling complex by transcriptional activators. Genes Dev. 13, 2369–2374 (1999).

23. Neely, K. E., Hassan, A. H., Brown, C. E., Howe, L. & Workman, J. L. Transcription activator interactions with multiple SWI/SNF subunits. Mol. Cell. Biol. 22, 1615–1625 (2002).

24. Papillon, J. P. N. et al. Discovery of Orally Active Inhibitors of Brahma Homolog (BRM)/SMARCA2 ATPase Activity for the Treatment of Brahma Related Gene 1 (BRG1)/SMARCA4-Mutant Cancers. J. Med. Chem. 61, 10155–10172 (2018).

25. Martin, B. J. E. et al. Global identification of SWI/SNF targets reveals compensation by EP400. Cell (2023) doi:10.1016/j.cell.2023.10.006.

26. Kundu, T. K. et al. Activator-Dependent Transcription from Chromatin In Vitro Involving Targeted Histone Acetylation by p300. Mol. Cell 6, 551–561 (2000).

27. Alerasool, N., Leng, H., Lin, Z.-Y., Gingras, A.-C. & Taipale, M. Identification and functional characterization of transcriptional activators in human cells. Mol. Cell 82, 677–695.e7 (2022).

28. Rada-Iglesias, A. et al. A unique chromatin signature uncovers early developmental enhancers in humans. Nature 470, 279–283 (2011).

29. Brower-Toland, B. et al. Specific contributions of histone tails and their acetylation to the mechanical stability of nucleosomes. J. Mol. Biol. 346, 135–146 (2005).

30. Lasko, L. M. et al. Discovery of a selective catalytic p300/CBP inhibitor that targets lineage-specific tumours. Nature 550, 128–132 (2017).

31. Raj, A., Peskin, C. S., Tranchina, D., Vargas, D. Y. & Tyagi, S. Stochastic mRNA synthesis in mammalian cells. PLoS Biol. 4, e309 (2006).

32. Suter, D. M. et al. Mammalian genes are transcribed with widely different bursting kinetics. Science 332, 472–474 (2011).

33. Chong, S., Chen, C., Ge, H. & Xie, X. S. Mechanism of transcriptional bursting in bacteria. Cell 158, 314–326 (2014).

34. Xiao, J. Y., Hafner, A. & Boettiger, A. N. How subtle changes in 3D structure can create large changes in transcription. Elife 10, (2021).

35. Zuin, J. et al. Nonlinear control of transcription through enhancer-promoter interactions. Nature 604, 571–577 (2022).

36. Gossen, M. & Bujard, H. Tight control of gene expression in mammalian cells by tetracycline-responsive promoters. Proc. Natl. Acad. Sci. U. S. A. 89, 5547–5551 (1992).

37. Kessler, D. S., Veals, S. A., Fu, X. Y. & Levy, D. E. Interferon-alpha regulates nuclear translocation and DNA-binding affinity of ISGF3, a multimeric transcriptional activator. Genes Dev. 4, 1753–1765 (1990).

38. Lazear, H. M., Schoggins, J. W. & Diamond, M. S. Shared and Distinct Functions of Type I and Type III Interferons. Immunity 50, 907–923 (2019).

39. Platanitis, E. et al. A molecular switch from STAT2-IRF9 to ISGF3 underlies interferon-induced gene transcription. Nat. Commun. 10, 2921 (2019).

40. Rengachari, S. et al. Structural basis of STAT2 recognition by IRF9 reveals molecular insights into ISGF3 function. Proc. Natl. Acad. Sci. U. S. A. 115, E601–E609 (2018).

41. Bluyssen, H. R. & Levy, D. E. Stat2 Is a Transcriptional Activator That Requires Sequence-specific Contacts Provided by Stat1 and p48 for Stable Interaction with DNA*. J. Biol. Chem. 272, 4600–4605 (1997).

42. Cui, K. et al. The chromatin-remodeling BAF complex mediates cellular antiviral activities by promoter priming. Mol. Cell. Biol. 24, 4476–4486 (2004).

43. Patel, M. C. et al. BRD4 coordinates recruitment of pause release factor P-TEFb and the pausing complex NELF/DSIF to regulate transcription elongation of interferon-stimulated genes. Mol. Cell. Biol. 33, 2497–2507 (2013).

44. Manry, J. et al. Evolutionary genetic dissection of human interferons. J. Exp. Med. 208, 2747–2759 (2011).

45. Krause, C. D. & Pestka, S. Cut, copy, move, delete: The study of human interferon genes reveal multiple mechanisms underlying their evolution in amniotes. Cytokine 76, 480–495 (2015).

46. Arimoto, K.-I., Miyauchi, S., Stoner, S. A., Fan, J.-B. & Zhang, D.-E. Negative regulation of type I IFN signaling. J. Leukoc. Biol. (2018) doi:10.1002/JLB.2MIR0817-342R.

47. Mostafavi, S. et al. Parsing the Interferon Transcriptional Network and Its Disease Associations. Cell 164, 564–578 (2016).

48. Cusanovich, D. A., Pavlovic, B., Pritchard, J. K. & Gilad, Y. The functional consequences of variation in transcription factor binding. PLoS Genet. 10, e1004226 (2014).

49. Lee, D. Y., Hayes, J. J., Pruss, D. & Wolffe, A. P. A positive role for histone acetylation in transcription factor access to nucleosomal DNA. Cell 72, 73–84 (1993).

50. Struhl, K. Histone acetylation and transcriptional regulatory mechanisms. Genes Dev. 12, 599–606 (1998).

51. Narita, T. et al. Enhancers are activated by p300/CBP activity-dependent PIC assembly, RNAPII recruitment, and pause release. Mol. Cell 81, 2166–2182.e6 (2021).

52. 52. Ferrie, J. J., et al. p300 Is an Obligate Integrator of Combinatorial Transcription Factor Inputs. *bioRxiv* (2023) doi:10.1101/2023.05.18.541220.

53. DelRosso, N. et al. Large-scale mapping and mutagenesis of human transcriptional effector domains. Nature 616, 365–372 (2023).

54. Durrant, M. G. et al. Systematic discovery of recombinases for efficient integration of large DNA sequences into the human genome. Nat. Biotechnol. 41, 488–499 (2023).

55. Tycko, J. et al. High-Throughput Discovery and Characterization of Human Transcriptional Effectors. Cell 183, 2020–2035.e16 (2020).

56. Iurlaro, M. et al. Mammalian SWI/SNF continuously restores local accessibility to chromatin. Nat. Genet. 53, 279–287 (2021).

57. Weinert, B. T. et al. Time-Resolved Analysis Reveals Rapid Dynamics and Broad Scope of the CBP/p300 Acetylome. Cell 174, 231–244.e12 (2018).

58. Teague, B. Cytoflow: A Python Toolbox for Flow Cytometry. *bioRxiv* 2022.07.22.501078 (2022) doi:10.1101/2022.07.22.501078.

59. Pedersen, B. S., Eyring, K., De, S., Yang, I. V. & Schwartz, D. A. Fast and accurate alignment of long bisulfite-seq reads. arXiv [q-bio.GN] (2014).

60. Virtanen, P. et al. SciPy 1.0: fundamental algorithms for scientific computing in Python. Nat. Methods 17, 261–272 (2020).

61. Corces, M. R. et al. An improved ATAC-seq protocol reduces background and enables interrogation of frozen tissues. Nat. Methods 14, 959–962 (2017).

62. Robinson, J. T. et al. Integrative genomics viewer. Nat. Biotechnol. 29, 24–26 (2011).

63. Schep, A. N., Wu, B., Buenrostro, J. D. & Greenleaf, W. J. chromVAR: inferring transcription-factor-associated accessibility from single-cell epigenomic data. Nat. Methods 14, 975–978 (2017).

64. Love, M. I., Huber, W. & Anders, S. Moderated estimation of fold change and dispersion for RNA-seq data with DESeq2. Genome Biol. 15, 550 (2014).

65. Marinov, G. K. ChIP-seq for the Identification of Functional Elements in the Human Genome. Methods Mol. Biol. 1543, 3–18 (2017).

66. Langmead, B., Trapnell, C., Pop, M. & Salzberg, S. L. Ultrafast and memory-efficient alignment of short DNA sequences to the human genome. Genome Biol. 10, R25 (2009).

67. Feng, J., Liu, T., Qin, B., Zhang, Y. & Liu, X. S. Identifying ChIP-seq enrichment using MACS. Nat. Protoc. 7, 1728–1740 (2012).

68. Marinov, G. K. Identification of Candidate Functional Elements in the Genome from ChIP-seq Data. Methods Mol. Biol. 1543, 19–43 (2017).

