## Supplemental Code for "Single-molecule chromatin configurations link transcription factor binding to expression in human cells": IFN_ATAC_chraccr.rmd

````
---
title: "R Notebook"
output: html_notebook
---


```{r}
# script modified from Sidney V.
# Import Libraries
suppressWarnings({
  library(ChrAccR)
  library(muLogR)
  library(muRtools)
  library(ggplot2)
  library(chromVAR)
  library(motifmatchr)
  library(JASPAR2018)
  library(BSgenome.Hsapiens.UCSC.hg38)
  library(dplyr)
  library(DESeq2)
  #library(biomaRt)
  library(GenomicFeatures)
  library(ChrAccRAnnotationHg38)
  library(tidyverse)
  library(ggpubr)
  library(ggrastr)
  library(ggrepel)
  library(viridis)
  library(Gviz)
  library(TFBSTools)
  library(reshape2)
})

source("/oak/stanford/groups/wjg/jschaepe/templates/R_scripts/general_funcs.R")
source("/oak/stanford/groups/wjg/jschaepe/templates/R_scripts/tf_funcs.R")

setwd("/oak/stanford/groups/wjg/jschaepe/projects/atac/20220921_julia_IFN/23.08.08_IFN_ATAC_dataset")


# Make dsa bam object -----------------------------------------------------
sampleAnnot <- read.csv("../sample_sheet.csv")


setConfigElement("annotationColumns", c("hours"))
setConfigElement("filteringCovgCount", 1L)
setConfigElement("filteringCovgReqSamples", 0.25)
setConfigElement("filteringSexChroms", TRUE)
setConfigElement("normalizationMethod", "quantile")

consensuspeaks <- getPeakSet.snakeATAC(sampleAnnot, filePrefixCol = "peakFilename", genome = "hg38", dataDir = "", sampleIdCol = "sampleName")
dsa_fromBam <- DsATAC.bam(sampleAnnot, "bamFilenameFull", "hg38", regionSets = list("consensuspeaks" = consensuspeaks), sampleIdCol="sampleName", 
                          pairedEnd=FALSE)
saveDsAcc(dsa_fromBam, "dsa_fromBam_v2")


# Add promoter region to the dsa object -----------------------------------
promoGr <- ChrAccRAnnotationHg38::getGeneAnnotation(type="promoterGr") # gets promotor regions
# Adds another region type (promoter)
dsa <- regionAggregation(dsa_fromBam, promoGr, "promoter", signal="insertions") 

saveDsAcc(dsa, "dsa_w_promoter_v2")


#plot(getCoord(dsa,"filteredConsensus")$score)

# Create ChrAccR Reports --------------------------------------------------

reportDir <- file.path(".", "ChrAccR_reports_v2")
# create the report (takes ~10 min on the example dataset)
createReport_summary(dsa, reportDir)

dsa_norm <- transformCounts(dsa, method = "RPKM")
dsa_norm <- transformCounts(dsa_norm, method = "log2")
dsa_norm <- transformCounts(dsa_norm, method = "quantile")
saveDsAcc(dsa_norm, "dsa_rpkmlog2qnorm_v2")

# create the report (takes ~1 min on the example dataset)
setConfigElement("chromVarRegionTypes", c("consensuspeaks","promoter"))
createReport_exploratory(dsa_norm, reportDir)

setConfigElement("regionTypes",c("consensuspeaks"))
setConfigElement("differentialColumns", c("hours"))
createReport_differential(dsa, reportDir)


# PCA ---------------------------------------------------------------------
dsa_norm <- loadDsAcc('dsa_rpkmlog2qnorm_v2/')
cm <- as.data.frame(dsa_norm@counts$consensuspeaks)
pcaCoord <- muRtools::getDimRedCoords.pca(t(cm))
annot <- getSampleAnnot(dsa_norm)

pdf("plots/PCA.pdf", width=9, height=7)
muRtools::getDimRedPlot(pcaCoord, annot=annot, colorCol="hours",ptSize=5) + 
                theme_classic() + theme(text=element_text(size=20),panel.border = element_rect(color ="black",fill = NA, size = 1))
dev.off()

# TF Motif Enrichment on differential peaks -------------------------------
dds <- readRDS("ChrAccR_reports_v2/differential_data/diffObj_consensuspeaks.rds") 

group1 <- "Cont" # reference level
contrast.col <- "hours"
region <- "consensuspeaks"
p.cutoff <- 0.05
log2fc.cutoff <- 0.58
  
## motif matching to JASPAR ------------------------------------------------
#this only needs to be run once per work session
motifs <- getMotifs() # this gets JASPAR2018 motif
genome <- "BSgenome.Hsapiens.UCSC.hg38"
Peaks <- as.data.frame(dsa_norm@coord[[region]],row.names = NULL, stringsAsFactors=FALSE) %>% 
  dplyr::select(c("seqnames","start","end"))
Peaks.Ranges <- makeGRangesFromDataFrame(Peaks, ignore.strand = T)
motif.scores <- matchMotifs(motifs, Peaks.Ranges, genome = genome, out="scores")
saveRDS(motif.scores, "dsa_norm_v2_motif.scores.rds")
motif.scores <- readRDS("dsa_norm_v2_motif.scores.rds")

for (group2 in c('1 hour', '3 hours', '6 hours', '12 hours', '24 hours')){
  message(group2)
  tf_motif_de_analysis(dds, dsa_norm, motifs, motif.scores, Peaks, 
                       '0 hours', group2, contrast.col, region, p.cutoff=p.cutoff, log2fc.cutoff=log2fc.cutoff) # function in tf_funcs.R
}
for (group2 in c('24 hours')){
  message(group2)
  tf_motif_de_analysis(dds, dsa_norm, motifs, motif.scores, Peaks, 
                       '0 hours', group2, contrast.col, region, p.cutoff=p.cutoff, log2fc.cutoff=log2fc.cutoff) # function in tf_funcs.R
}
```
````
