## Supplemental Code for "Single-molecule chromatin configurations link transcription factor binding to expression in human cells": IFN_ATAC_footprinting.r

```
library(ChrAccR)
library(tidyverse)

setwd("/oak/stanford/groups/wjg/jschaepe/projects/atac/20220921_julia_IFN/23.08.08_IFN_ATAC_dataset")

dsa <- loadDsAcc("dsa_w_promoter_v2/")
dsa_cpm <- transformCounts(dsa, method = "CPM")

# read in Vierstra v2 archetype motifs
if (!dir.exists("footprinting")){
  dir.create("footprinting")
}

# browse motifs with https://resources.altius.org/~jvierstra/projects/motif-clustering-v2.1beta/ 
url <- "https://jeffgranja.s3.amazonaws.com/ArchR/Annotations/Vierstra_Archetype_Motifs_v2.1.rds"
annoPath <- "footprinting"
motifFile <- file.path(annoPath, basename(url))

if (!file.exists(motifFile)){
  download.file(
    url = url, 
    destfile = file.path(annoPath, basename(url)),
    quiet = FALSE
  )
}
vierstra.motifs <- readRDS(motifFile)

motifNames <- grep("(AC0188)", names(vierstra.motifs), value=TRUE) # searching for patterns
samples <- getSampleAnnot(dsa_cpm)$sampleName
fps <- getMotifFootprints(dsa_cpm, motifNames, samples, motifDb=vierstra.motifs) # takes 5 hours

saveRDS(fps, "footprinting/fps_vierstra_ISRE_v2.rds")

fps <- readRDS("footprinting/fps_vierstra_ISRE_v2.rds")

plot_motifs_smooth <- function(fps, motif, dsa){
  fp <- fps[[motif]]$footprintDf
  meta <- dsa@sampleAnnot
  fp$hours <- meta[fp$sampleId,"hours"]
  fp <- fp %>%
    mutate(hours_int = as.integer(str_extract(hours, "\\S+")))
  fp <- fp %>%
    mutate(hours_str = as.character(hours_int))
  fp <- fp[order(fp$hours_int, decreasing = FALSE), ]
  tmp <- fp[fp$hours=="0 hours",]  %>% group_by(pos) %>% summarise(avg_ctl_countNormBiasCor = mean(countNormBiasCor), 
                                                                   avg_ctl_countNorm = mean(countNorm),
                                                                   avg_ctl_Tn5biasNorm = mean(Tn5biasNorm)) %>% as.data.frame
  fp <- fp %>% dplyr::mutate(avg_ctl_countNormBiasCor = lapply(fp$pos, function(p){tmp[tmp$pos==p, "avg_ctl_countNormBiasCor"]}) %>% unlist,
                             avg_ctl_countNorm = lapply(fp$pos, function(p){tmp[tmp$pos==p, "avg_ctl_countNorm"]}) %>% unlist)
  
  fp <- fp %>% group_by(hours_int,hours_str, hours, pos) %>% summarise(countNorm = mean(countNorm),
                                                                       countNormBiasCor = mean(countNormBiasCor),
                                                                       countNormBiasCor_ctlNorm = mean(countNormBiasCor / avg_ctl_countNormBiasCor),
                                                                       countNorm_ctlNorm = mean(countNorm / avg_ctl_countNorm),
                                                                       Tn5biasNorm = mean(Tn5biasNorm))
  
  # clean up motif name for file name
  motif_out <- gsub("[[:punct:]]", "_", motif)

  ggplot(fp) + aes(x=pos, y=countNorm, col=hours) + geom_smooth(method = "loess", span=0.1, se = FALSE)  + scale_color_manual(values = c('#f8cbcd','#f2989b','#961b23','#050505','#ed676a','#ea3840')) + theme_classic()
  ggsave(paste0("footprinting/",motif_out,"_countNorm_smooth.pdf"), width=6, height=6)

  ggplot(fp) + aes(x=pos, y=countNormBiasCor, col=hours) + geom_smooth(method = "loess", span=0.1, se = FALSE) + scale_color_manual(values = c('#f8cbcd','#f2989b','#961b23','#050505','#ed676a','#ea3840'))+ theme_classic()
  ggsave(paste0("footprinting/",motif_out, "_countNormBiasCor_smooth.pdf"), width=6, height=6)
  
  ggplot(fp) + aes(x=pos, y=countNormBiasCor_ctlNorm, col=hours) + geom_smooth(method = "loess", span=0.1, se = FALSE) + scale_color_manual(values = c('#f8cbcd','#f2989b','#961b23','#050505','#ed676a','#ea3840'))+ theme_classic()
  ggsave(paste0("footprinting/",motif_out, "_countNormBiasCor_ctlNorm_smooth.pdf"), width=6, height=6)
  
  ggplot(fp) + aes(x=pos, y=countNorm_ctlNorm, col=hours) + geom_smooth(method = "loess", span=0.1, se = FALSE) + scale_color_manual(values = c('#f8cbcd','#f2989b','#961b23','#050505','#ed676a','#ea3840'))+ theme_classic()
  ggsave(paste0("footprinting/",motif_out, "_countNorm_ctlNorm_smooth.pdf"), width=6, height=6)

  ggplot(fp) + aes(x=pos, y=Tn5biasNorm, col=hours) + geom_smooth(method = "loess", span=0.1, se = FALSE) + scale_color_manual(values = c('#f8cbcd','#f2989b','#961b23','#050505','#ed676a','#ea3840'))+ theme_classic()
  ggsave(paste0("footprinting/",motif_out, "_Tn5biasNorm_smooth.pdf"), width=6, height=6)
  invisible(dev.off())
}

plot_motifs_smooth(fps, "AC0188|IRF/STAT|IRF", dsa_cpm)
```
