## Supplemental Code for "Single-molecule chromatin configurations link transcription factor binding to expression in human cells": IFN_RNA_processing.r

```
# follow this tutorial:
# http://bioconductor.org/help/course-materials/2016/CSAMA/lab-3-rnaseq/rnaseq_gene_CSAMA2016.pdf

# if (!requireNamespace("BiocManager", quietly = TRUE))
#   install.packages("BiocManager")
# 
# BiocManager::install("GenomicFeatures")
# BiocManager::install("topGO")
# BiocManager::install("PCAtools")
# BiocManager::install("Homo.sapiens")

suppressWarnings({
  library('Rsamtools')
  library('GenomicFeatures')
  library('GenomicAlignments')
  library('BiocParallel')
  library('DESeq2')
  library('PCAtools')
  library('ggplot2')
  library('pheatmap')
  library('AnnotationDbi')
  library('Homo.sapiens')
  library('topGO') 
  library('RColorBrewer')
  library('ggrepel')
  library('readr')
  library('tibble')
  library('grid')
  library('lattice')
  library('gridExtra')
  library('patchwork')
})

source("/oak/stanford/groups/wjg/jschaepe/projects/rna/230810_IFN_RNAseq/RNA-seq_functions.R")
setwd("/oak/stanford/groups/wjg/jschaepe/projects/rna/230810_IFN_RNAseq")

# read in sample table that contains metadata (e.g. cell type, treatment, etc.)
sampleTable <- read.csv('20211004_sample_table.csv',row.names=1)

countsFile <- read.csv("se_counts.csv", stringsAsFactors=FALSE)
se <- SummarizedExperiment(data.matrix(countsFile))
colData(se) <- DataFrame(sampleTable)
se
assay(se)

# filter out any genes that have fewer than 5 counts
se2 <- se[rowSums(assay(se)) >=5, ]

# The rows of the samples (K562 WT +/- IFN) are 21-24
dds <- get_dds(c(21:24), 'IFN', 'Rep')
plot <- make_volcano_plot(c(21:24), 'IFN', 'Rep', 'IFN', 'noIFN')
ggsave('volcano.pdf', plot=plot, width=6, height=6)
```
