## Supplemental Code for "Single-molecule chromatin configurations link transcription factor binding to expression in human cells": isg_preaccessibility_and_motifs.ipynb

```
# %% [code]
import pandas as pd
import numpy as np
from matplotlib import pyplot as plt
import seaborn as sns
from matplotlib import pylab
import seaborn as sns
import os
from os import path
from scipy.stats import pearsonr, spearmanr, chi2_contingency
from scipy.cluster.hierarchy import fcluster, fclusterdata, linkage, dendrogram 
import matplotlib.patches as patches
from copy import deepcopy
import pickle

# common formatting
plt.style.use('../../synthetic-enhancer-footprinting.mplstyle')

import sys  
# sys.path.insert(0, '/oak/stanford/groups/wjg/bgrd/papers/synthetic_enhancer_footprinting/code/amplicon-smf/workflow/scripts')
sys.path.insert(0, '/oak/stanford/groups/wjg/bgrd/papers/synthetic_enhancer_footprinting/data/partition_function_model_outputs/231114_first_pass_at_reunning_everything/amplicon-smf/workflow/scripts')

import common, fit_partition_function_model_v3


import importlib
importlib.reload(common)
importlib.reload(fit_partition_function_model_v3)
%matplotlib inline

import matplotlib.font_manager

output_path = '/oak/stanford/groups/wjg/jschaepe/projects/smf/synthetic-enhancer-footprinting-outputs/'

fontsize=6
markersize=20
legend_fontsize = 5

import gzip


# %% [code]
# here is where the atac processing lives: $OAK/jschaepe/projects/atac/20220921_julia_IFN/23.08.08_IFN_ATAC_dataset
# the snake atac output is in 20220921_julia_IFN/snakeATAC_sing
# rna data and processing lives here: $OAK/jschaepe/projects/rna/230810_IFN_RNAseq
# motifs from georgi: /oak/stanford/groups/akundaje/marinovg/motifs/FIMO-CIS-BP-Homo_sapiens.hg20.fimo.txt.annotated.bgz
# greenleaf tss: /oak/stanford/groups/wjg/share/resources/hg38/hg38.tss.bed
# engreitz tss: /oak/stanford/groups/wjg/bgrd/data/genomes/hg38/CollapsedGeneBounds.hg38.TSS500bp.bed
# bash commands live here: /oak/stanford/groups/wjg/bgrd/papers/synthetic_enhancer_footprinting/data/ifn_analyses/
basedir = '/oak/stanford/groups/wjg/bgrd/papers/synthetic_enhancer_footprinting/data/ifn_analyses/231219_ifn_analyses'

# %% [code]
# first step is to make confident TSS annotations for ISGs
de_genes_file = '/oak/stanford/groups/wjg/jschaepe/projects/rna/230810_IFN_RNAseq/DEseq_results.csv'
de_genes = pd.read_csv(de_genes_file, index_col=0)

# select just the ones that go up
p_thresh = 0.05
l2fc_thresh = 1

isgs = de_genes.loc[(de_genes.padj < p_thresh)&(de_genes.log2FoldChange > l2fc_thresh)]
isg_ids = isgs.index.tolist()
print(len(isg_ids))
isgs.head()

# %% [code]
# one way is to load hg38 tss file from wjg atac pipeline
# tss
wjg_tss_file = '/oak/stanford/groups/wjg/share/resources/hg38/hg38.tss.bed'
wjg_tss = pd.read_table(wjg_tss_file, header=None)
isg_tss = wjg_tss.loc[wjg_tss[5].isin(isgs.symbol)]
isg_tss.to_csv(path.join(basedir, 'isg_tss.wjg.bed'), sep='\t', header=False, index=False)
print(len(isg_tss))
isg_tss.head()

# %% [code]
# another way is to load the engreitz annotation, and try joining either on ensembl or symbol
abc_tss_file = '/oak/stanford/groups/wjg/bgrd/data/genomes/hg38/CollapsedGeneBounds.hg38.TSS500bp.bed'
abc_tss = pd.read_table(abc_tss_file)
isg_tss_symbol = abc_tss.loc[abc_tss['name'].isin(isgs.symbol)]
isg_tss_ensembl = abc_tss.loc[abc_tss['Ensembl_ID'].isin(isg_ids)]
isg_tss_symbol.to_csv(path.join(basedir, 'isg_tss.engreitz_symbol.bed'), sep='\t', header=False, index=False)
isg_tss_ensembl.to_csv(path.join(basedir, 'isg_tss.engreitz_ensembl.bed'), sep='\t', header=False, index=False)
print(len(isg_tss_symbol))
print(len(isg_tss_ensembl))
isg_tss_symbol.head()

# %% [code]
# get null genes
f = path.join(basedir, '20221129_normalized_tpm_counts.csv')
tpm = pd.read_csv(f, index_col=0)
tpm['no_ifn_tpm'] = tpm[['K562_WT_noTX_BR1_S5','K562_WT_noTX_BR2_S6']].mean(axis=1)
tpm.head()

l2fc_thresh_null = 0.5

no_change_genes = de_genes.loc[de_genes.log2FoldChange.between(-1*l2fc_thresh_null,l2fc_thresh_null)].index.tolist()

null_genes = []

for isg in isg_ids:
    basemean = tpm.loc[isg,'no_ifn_tpm']
    null_gene = np.abs(tpm.loc[no_change_genes,'no_ifn_tpm'] - basemean).sort_values().index[0]
    null_genes.append(null_gene)
    
null_tss_ensembl = abc_tss.loc[abc_tss['Ensembl_ID'].isin(null_genes)].copy()
null_tss_ensembl.to_csv(path.join(basedir, 'null_tss.engreitz_ensembl.bed'), sep='\t', header=False, index=False)

# %% [code]
# now go do stuff in the notebook

# %% [code]
motifs = ['STAT1', 'STAT2', 'IRF9']

dfs = []

for motif in motifs:
    f = path.join(basedir, 'isg_tss.engreitz_ensembl.{}.counts'.format(motif))
    df = pd.read_table(f, header=None, names=['chr','start','end','gene','.','strand','id','type',motif])
    dfs.append(df)

isg_df = dfs[0].merge(dfs[1], on='gene', suffixes=['','_1']).merge(dfs[2], on='gene', suffixes=['','_2'])[['chr','start','end','gene','.','strand','id','type']+motifs]
isg_df['motif_sum'] = isg_df[motifs].sum(axis=1)
isg_df['motif_max'] = isg_df[motifs].max(axis=1)
isg_df['stat2_irf9'] = isg_df[['STAT2', 'IRF9']].max(axis=1)

# now do for null
null_dfs = []

for motif in motifs:
    f = path.join(basedir, 'null_tss.engreitz_ensembl.{}.counts'.format(motif))
    df = pd.read_table(f, header=None, names=['chr','start','end','gene','.','strand','id','type',motif])
    null_dfs.append(df)

null_df = null_dfs[0].merge(null_dfs[1], on='gene', suffixes=['','_1']).merge(null_dfs[2], on='gene', suffixes=['','_2'])[['chr','start','end','gene','.','strand','id','type']+motifs]
null_df['motif_sum'] = null_df[motifs].sum(axis=1)
null_df['motif_max'] = null_df[motifs].max(axis=1)
null_df['stat2_irf9'] = null_df[['STAT2', 'IRF9']].max(axis=1)

isg_df.head(), null_df.head()

# %% [code]
fig, ax = plt.subplots(figsize=(4/2.5,3/2.5))

isg_color = '#F6999B'
null_color = '0.5'
sixhr_color = '#d51317'

col = 'stat2_irf9'

df = pd.DataFrame({'ISG': isg_df[col].value_counts(normalize=True), 'Background': null_df[col].value_counts(normalize=True)})

df.plot.bar(ax=ax, rot=0, color = [isg_color, null_color])

ax.set_xlabel('Number ISRE in promoter', fontsize = fontsize)
ax.set_xticks([0,1,2,3,4])
ax.set_xticklabels([0,1,2,3,4], fontsize = fontsize)
ax.set_ylabel('Fractions promoters', fontsize = fontsize)
# ax.set_ylim(0.05,0.55)
ax.set_yticks([0.0, 0.2, 0.4, 0.6, 0.8])
ax.set_yticklabels([0.0, 0.2, 0.4, 0.6, 0.8], fontsize = fontsize)
ax.tick_params(axis='both', labelsize=fontsize)
ax.set_box_aspect(0.8)
ax.spines['top'].set_visible(False)
ax.spines['right'].set_visible(False)
# ax.get_legend().set_visible(False)
plt.legend(fontsize=legend_fontsize, markerscale=0.5, frameon=False, title='', title_fontsize=legend_fontsize, bbox_to_anchor=(1,1))

plt.savefig('{}/fig4_plots/fig4_supplement/isre_matches_at_isgs.pdf'.format(output_path), dpi=300)


# %% [code]
motif_thresh = 2

bonafide_isgs = isg_df.loc[isg_df[col] > motif_thresh,'gene'].tolist()
bonafide_isgs_ids = isg_df.loc[isg_df[col] > motif_thresh,'id'].tolist()

bonafide_null_genes = []

null_to_isg_map = {}

for isg in bonafide_isgs_ids:
    basemean = tpm.loc[isg,'no_ifn_tpm']
    null_gene = np.abs(tpm.loc[no_change_genes,'no_ifn_tpm'] - basemean).sort_values().index[0]
    bonafide_null_genes.append(null_gene)
    null_to_isg_map[null_gene] = isg


# %% [code]
samples = ['JMS-0IFN-b1','JMS-0IFN-b2',]
samples_6hr = ['JMS-6IFN-b1','JMS-6IFN-b2']

read_depths = {'JMS-0IFN-b1': 16539520, 
               'JMS-0IFN-b2': 19947866, 
               'JMS-6IFN-b1': 24661726, 
               'JMS-6IFN-b2': 21147274}

dfs = []

for samp in samples:
    f = path.join(basedir, '{}.isg_tss.engreitz_ensembl.counts'.format(samp))
    df = pd.read_table(f, header=None, names=['chr','start','end','gene','.','strand','id','type','pos',samp])
    df['{}_rpm'.format(samp)] = df[samp] / read_depths[samp] * 1e6
    dfs.append(df)

isg_df = dfs[0].merge(dfs[1], on=['gene','pos'], suffixes=['','_1'])[['chr','start','end','gene','.','strand','id','type','pos']+samples+[s+'_rpm' for s in samples]]
isg_df['rpm'] = isg_df.filter(like='rpm').mean(axis=1)

# now for null
null_dfs = []

for samp in samples:
    f = path.join(basedir, '{}.null_tss.engreitz_ensembl.counts'.format(samp))
    df = pd.read_table(f, header=None, names=['chr','start','end','gene','.','strand','id','type','pos',samp])
    df['{}_rpm'.format(samp)] = df[samp] / read_depths[samp] * 1e6
    null_dfs.append(df)

null_df = null_dfs[0].merge(null_dfs[1], on=['gene','pos'], suffixes=['','_1'])[['chr','start','end','gene','.','strand','id','type','pos']+samples+[s+'_rpm' for s in samples]]
null_df['rpm'] = null_df.filter(like='rpm').mean(axis=1)

# now do 6hr IFN
sixhr_dfs = []

for samp in samples_6hr:
    f = path.join(basedir, '{}.isg_tss.engreitz_ensembl.counts'.format(samp))
    df = pd.read_table(f, header=None, names=['chr','start','end','gene','.','strand','id','type','pos',samp])
    df['{}_rpm'.format(samp)] = df[samp] / read_depths[samp] * 1e6
    sixhr_dfs.append(df)

sixhr_df = sixhr_dfs[0].merge(sixhr_dfs[1], on=['gene','pos'], suffixes=['','_1'])[['chr','start','end','gene','.','strand','id','type','pos']+samples_6hr+[s+'_rpm' for s in samples_6hr]]
sixhr_df['rpm'] = sixhr_df.filter(like='rpm').mean(axis=1)

isg_df

# %% [code]
isg_df_to_use = isg_df.loc[isg_df.gene.isin(bonafide_isgs)]
null_df_to_use = null_df.loc[null_df.id.isin(bonafide_null_genes)]
sixhr_df_to_use = sixhr_df.loc[sixhr_df.gene.isin(bonafide_isgs)]

isg_mean = isg_df_to_use.groupby('pos')['rpm'].mean()
isg_sem = isg_df_to_use.groupby('pos')['rpm'].sem()
null_mean = null_df_to_use.groupby('pos')['rpm'].mean()
null_sem = null_df_to_use.groupby('pos')['rpm'].sem()
sixhr_mean = sixhr_df_to_use.groupby('pos')['rpm'].mean()
sixhr_sem = sixhr_df_to_use.groupby('pos')['rpm'].sem()

window = 10

isg_mean_smoothed = np.convolve(isg_mean, np.ones(window)/window, mode='valid')
null_mean_smoothed = np.convolve(null_mean, np.ones(window)/window, mode='valid')
sixhr_mean_smoothed = np.convolve(sixhr_mean, np.ones(window)/window, mode='valid')
isg_sem_smoothed = np.convolve(isg_sem, np.ones(window)/window, mode='valid')
null_sem_smoothed = np.convolve(null_sem, np.ones(window)/window, mode='valid')
sixhr_sem_smoothed = np.convolve(sixhr_sem, np.ones(window)/window, mode='valid')

fig, ax = plt.subplots(figsize=(4/2.5,3/2.5))

ax.fill_between(range(500-window+1), null_mean_smoothed-null_sem_smoothed,null_mean_smoothed+null_sem_smoothed, alpha=0.2, color=null_color)
ax.fill_between(range(500-window+1), sixhr_mean_smoothed-sixhr_sem_smoothed,sixhr_mean_smoothed+sixhr_sem_smoothed, alpha=0.2, color=sixhr_color)
ax.fill_between(range(500-window+1), isg_mean_smoothed-isg_sem_smoothed,isg_mean_smoothed+isg_sem_smoothed, alpha=0.2, color=isg_color)
ax.plot(null_mean_smoothed, label='Background', c=null_color)
ax.plot(sixhr_mean_smoothed, label='+IFN', c=sixhr_color)
ax.plot(isg_mean_smoothed, label='ISG', c=isg_color)

ax.set_xlabel('Position relative to TSS', fontsize = fontsize)
ax.set_xticks([0,250,500])
ax.set_xticklabels([-250,0,250], fontsize = fontsize)
ax.set_ylabel('ATAC-seq RPM', fontsize = fontsize)
ax.set_ylim(0.05,0.9)
ax.set_yticks([0.2, 0.4, 0.6, 0.8])
ax.set_yticklabels([0.2, 0.4, 0.6, 0.8], fontsize = fontsize)
ax.tick_params(axis='both', labelsize=fontsize)
ax.set_box_aspect(0.8)
ax.spines['top'].set_visible(False)
ax.spines['right'].set_visible(False)
# ax.get_legend().set_visible(False)
plt.legend(fontsize=legend_fontsize, markerscale=0.5, frameon=False, title='', title_fontsize=legend_fontsize, bbox_to_anchor=(1,1))

plt.savefig('{}/fig4_plots/isg_preaccessibility_by_atac.with_6hr.pdf'.format(output_path), dpi=300)


# %% [code]
```
