## Supplemental Code for "Single-molecule chromatin configurations link transcription factor binding to expression in human cells": RNA-seq_functions.r

```
getwd()

# follow this tutorial:
# http://bioconductor.org/help/course-materials/2016/CSAMA/lab-3-rnaseq/rnaseq_gene_CSAMA2016.pdf

# if (!requireNamespace("BiocManager", quietly = TRUE))
#   install.packages("BiocManager")
# 
# BiocManager::install("GenomicFeatures")

library('Rsamtools')
library('GenomicFeatures')
library('GenomicAlignments')
library('BiocParallel')
library('DESeq2')
library('PCAtools')
library('ggplot2')
library('pheatmap')
library('AnnotationDbi')
library('Homo.sapiens')
library('topGO')
library('RColorBrewer')
library('ggrepel')
library('readr')
library('tibble')
library('grid')
library('lattice')
library('gridExtra')
library('patchwork')
#Code written by Connor Ludwig, 2021 August
#Adapted for use by Abby Thurm, 2021 October

###TO USE ALL OF THE FOLLOWING FUNCTIONS:####

#put this code at the top of your new R notebook to access functions:
#source("F:/Connor/20210900_RNASeq_HTinduce/RNA-seq_functions.R")
#or copy the script into your new directory and call source("RNA-seq_functions.R") directly
#functions can then be called and used as described below directly in your new console

#rows: rows of your sample table you'd like to use, that correspond to samples you want to
#differentially analyze
#eg: c(1:4, 7:12); or just 1:4

#var: property you want to var, passed in as string directly as in sample table
#eg: "dox", "IFN"

#control: property you're controlling, passed in as string directly as in sample table
#eg: "Rep"

#treat: experimental condition of your variable of interest, passed in as string
#exactly as in your filenames/sample table
#eg: "dox-48h", "dox-12h", "IFN", "pCL073"

#notreat: control condition of your variable, passed in as string exactly as in your 
#sample table
#eg: "nodox", "noIFN"

#filename: name the file of csv (as string) the function will give you (DESEq, GO terms, etc)
#path name not necessary - will output to same directory as files
#eg: "mygoterms.csv"

#you do not need to explicitly make a dds, just choose if you want to make PCA/eigencor/DESEq/etc
#**********
get_dds <- function(rows, var, control) {
  se1 <- se2[,rows]
  design <- sprintf('~%s + %s', control, var)
  dds <- DESeqDataSet(se1, design = as.formula(design))
  return(dds)
}

#make PCA plot, plot loadings
make_PCA_plots <- function(rows, var, control){
  se1 <- se2[,rows]
  dds <- get_dds(rows, var, control)
  vsd <- vst(dds)
  
  p <- pca(assay(vsd), metadata = colData(se1))
  
  plot1 <- biplot(p, x='PC1', y='PC2', colby=var, shape=control,
                  legendPosition='right',
                  title='K562 HT-Induce RNA-seq PCA')
  plot2 <- plotloadings(p,
                        components=getComponents(p, c(1,2,3,4)),
                        rangeRetain = 0.01,
                        labSize = 3.0,
                        title = 'Loadings plot',
                        subtitle = 'PC1, PC2, PC3, PC4',
                        caption = 'Top 1% variables',
                        shape = 16,
                        shapeSizeRange = c(3,3),
                        col = c('limegreen', 'black', 'red3'),
                        drawConnectors = TRUE)
  plot1+plot2
}

#eigencorplot only relevant if you want to analyze all of your samples and look at 
#PCAs across entire dataset
make_eigencorplot <- function(rows, var, control) {
  se1 <- se2[,rows]
  dds <- get_dds(rows, var, control)
  vsd <- vst(dds)
  
  p <- pca(assay(vsd), metadata = colData(se1))
  #dds <- DESeqDataSet(se2, design = ~ Rep + IFN)
  #vsd <- vst(dds)
  # plot <- eigencorplot(p,metavars = c('plasmid', 'IFN', 'dox', 'Rep'), col=brewer.pal(n = 9, name = "RdBu"))
  plot <- eigencorplot(p,metavars = c('plasmid', 'dox', 'Rep'), col=brewer.pal(n = 9, name = "RdBu"))
  return(plot)
}

#get initial DESeq analysis of up/downregulated genes
get_DESeq <- function(rows, var, control, treat, notreat){
  dds <- get_dds(rows, var, control)
  vsd <- vst(dds)
  dds2 <- DESeq(dds)
  res2 <- results(dds2, alpha=0.05, lfcThreshold = 0, contrast = c(var, treat, notreat))
  res <- res2[ !is.na(res2$padj), ]
  
  DESeq2::plotMA(res, ylim=c(-12,12))
}

#get heatmap of top 30 genes changes in your conditions
get_heatmap <- function(rows, var, control, treat, notreat){
  dds <- get_dds(rows, var, control)
  vsd <- vst(dds)
  dds2 <- DESeq(dds)
  
  res2 <- results(dds2, alpha=0.05, lfcThreshold = 0, contrast = c(var, treat, notreat))
  res <- res2[ !is.na(res2$padj), ]

  mat <- assay(vsd)[ head(order(res$padj),30), ]
  mat <- mat - rowMeans(mat)
  df <- as.data.frame(colData(vsd)[,c(control, var)])
  mat <- as.data.frame(mat)
  row <- mapIds(Homo.sapiens,
                keys=row.names(mat),
                column="SYMBOL",
                keytype="ENSEMBL",
                multiVals="first")
  rownames(mat) <- ifelse(!is.na(row), row, row.names(mat))
  mat <- t(as.matrix(mat))
  plot1 <- pheatmap(mat, annotation_row=df, angle_col = 45, labels_row = c('', '', '', '', '', '', '', '', '', '', '', ''))
  return(plot1)
}

#get top 30 genes in table of your conditions
get_ordered_results <- function(rows, var, control, treat, notreat, filename) {
  dds <- get_dds(rows, var, control)
  vsd <- vst(dds)
  dds2 <- DESeq(dds)
  
  res2 <- results(dds2, alpha=0.05, lfcThreshold = 0, contrast = c(var, treat, notreat))
  res <- res2[ !is.na(res2$padj), ]
  
  row <- mapIds(Homo.sapiens,
                keys=row.names(res),
                column="SYMBOL",
                keytype="ENSEMBL",
                multiVals="first")
  res$symbol <- ifelse(!is.na(row), row, row.names(res))
  
  resOrdered <- res[order(res$log2FoldChange),]
  resOrderedDf <- as.data.frame(resOrdered)
  # path = sprintf("%s_DESeq_%s_%s", date, var, control)
  write.csv(resOrderedDf, file = filename)
  return(resOrderedDf)
}

#get GO terms upregulated in your conditions
get_GO_up <- function(rows, var, control, treat, notreat, filename) {
  dds <- get_dds(rows, var, control)
  vsd <- vst(dds)
  dds2 <- DESeq(dds)
  
  res2 <- results(dds2, alpha=0.05, lfcThreshold = 0, contrast = c(var, treat, notreat))
  res <- res2[ !is.na(res2$padj), ]
  
  row <- mapIds(Homo.sapiens,
                keys=row.names(res),
                column="SYMBOL",
                keytype="ENSEMBL",
                multiVals="first")
  res$symbol <- ifelse(!is.na(row), row, row.names(res))
  
  genelistUp <-factor(as.integer(res$padj <0.05 & res$log2FoldChange > 1))
  names(genelistUp) <- rownames(res)
  
  myGOdata_Up <- new("topGOdata",
                     ontology = "BP",
                     allGenes = genelistUp,
                     nodeSize = 10,
                     annot = annFUN.org, mapping = "org.Hs.eg.db", ID = "ensembl")
  
  goTestResults <- runTest(myGOdata_Up, algorithm = "elim", statistic = "fisher")
  GenTableDF_Up <- GenTable(myGOdata_Up, goTestResults, topNodes=25)
  write.csv(GenTableDF_Up, file=filename)
  return(GenTableDF_Up)
}

#get GO terms downregulated in your conditions
get_GO_down <- function(rows, var, control, treat, notreat, filename) {
  dds <- get_dds(rows, var, control)
  vsd <- vst(dds)
  dds2 <- DESeq(dds)
  
  res2 <- results(dds2, alpha=0.05, lfcThreshold = 0, contrast = c(var, treat, notreat))
  res <- res2[ !is.na(res2$padj), ]
  
  row <- mapIds(Homo.sapiens,
                keys=row.names(res),
                column="SYMBOL",
                keytype="ENSEMBL",
                multiVals="first")
  res$symbol <- ifelse(!is.na(row), row, row.names(res))
  
  genelistDown <-factor(as.integer(res$padj <0.05 & res$log2FoldChange < -1))
  names(genelistDown) <- rownames(res)
  
  myGOdata_Down <- new("topGOdata",
                       ontology = "BP",
                       allGenes = genelistDown,
                       nodeSize = 10,
                       annot = annFUN.org, mapping = "org.Hs.eg.db", ID = "ensembl")
  
  goTestResults <- runTest(myGOdata_Down, algorithm = "elim", statistic = "fisher")
  GenTableDF_Down <- GenTable(myGOdata_Down, goTestResults, topNodes=25)
  write.csv(GenTableDF_Down, file=filename)
  return(GenTableDF_Down)
}

#make volcano plot across your conditions
make_volcano_plot <- function(rows, var, control, treat, notreat) {
  dds <- get_dds(rows, var, control)
  vsd <- vst(dds)
  dds2 <- DESeq(dds)
  
  res2 <- results(dds2, alpha=0.05, lfcThreshold = 0, contrast = c(var, treat, notreat))
  res <- res2[ !is.na(res2$padj), ]
  
  row <- mapIds(Homo.sapiens,
                keys=row.names(res),
                column="SYMBOL",
                keytype="ENSEMBL",
                multiVals="first")
  res$symbol <- ifelse(!is.na(row), row, row.names(res))
  
  low = -1
  high = 1
  pthresh = 5e-2
  res <- as.data.frame(res)
  res$diffexpressed <- 'NO'
  res$diffexpressed[res$log2FoldChange > high & res$pvalue < pthresh] <- 'UP'
  res$diffexpressed[res$log2FoldChange < low & res$pvalue < pthresh] <- 'DOWN'
  
  g <- ggplot(data=res, aes(x=log2FoldChange, y=-log10(pvalue), col=diffexpressed)) + geom_point() + theme_minimal()
  g2 <- g + geom_vline(xintercept=c(low, high), col="red") +
    geom_hline(yintercept=-log10(pthresh), col="red")
  mycolors <- c("blue", "red", "black")
  names(mycolors) <- c("DOWN", "UP", "NO")
  g3 <- g2 + scale_colour_manual(values = mycolors)
  
  res$delabel <- NA
  res$delabel[res$diffexpressed != "NO"] <- res$symbol[res$diffexpressed != "NO"]
  plot <- ggplot(data=res, aes(x=log2FoldChange, y=-log10(pvalue), col=diffexpressed, label=delabel)) +
    geom_point() + theme_minimal() + geom_text_repel() + scale_color_manual(values=c('blue', 'black', 'red')) +
    geom_vline(xintercept=c(low, high), col='red') + geom_hline(yintercept=-log10(pthresh), color='red')
  title = sprintf('%s vs %s: Volcano Plot', treat, notreat)
  plot<-plot+ggtitle(title)
  return(plot)
}
```
